# Variation and variability in *Drosophila* grooming behavior

**DOI:** 10.1101/2020.08.15.252627

**Authors:** Joshua M. Mueller, Neil Zhang, Jean M. Carlson, Julie H. Simpson

**Affiliations:** Interdepartmental Graduate Program in Dynamical Neuroscience, University of California, Santa Barbara; Department of Physics, University of California, Santa Barbara; Department of Molecular, Cellular, & Developmental Biology, University of California, Santa Barbara

**Author notes:** Experimental design: J.H.S. Data collection: J.H.S., N.Z.; data analysis: J.M.M.; manuscript writing: J.M.M.; manuscript editing: J.M.M., J.H.S., J.M.C. J.M.M. and J.H.S. contributed equally to this work.

## Abstract

Behavioral differences can be observed between species or populations (variation) or between individuals in a genetically similar population (variability). Here, we investigate genetic differences as a possible source of variation and variability in *Drosophila* grooming. Grooming confers survival and social benefits. Grooming features of five *Drosophila* species exposed to a dust irritant were analyzed. Aspects of grooming behavior, such as anterior to posterior progression, were conserved between and within species. However, significant differences in activity levels, proportion of time spent in different cleaning movements, and grooming syntax were identified between species. All species tested showed individual variability in the order and duration of action sequences. Genetic diversity was not found to correlate with grooming variability within a species: *melanogaster* flies bred to increase or decrease genetic heterogeneity exhibited similar variability in grooming syntax. Individual flies observed on consecutive days also showed grooming sequence variability. Standardization of sensory input using optogenetics reduced but did not eliminate this variability. In aggregate, these data suggest that sequence variability may be a conserved feature of grooming behavior itself. These results also demonstrate that large genetic differences result in distinguishable grooming phenotypes (variation), but that genetic heterogeneity within a population does not necessarily correspond to an increase in the range of grooming behavior (variability).

## Introduction

For over a century, researchers have understood that phenotypic variation arises from genotypic differences between organisms. Animal behavior contains phenotypes partially under genetic control, so a sizable research effort has been dedicated to uncovering specific genes associated with observable differences in behavior between and within species (1, 2). Different mouse species exhibit variation in monogamy and parental care, and different fly species show variation in courtship song and food preferences (3–6). From endangered species to agricultural crops to virus variants, genetic diversity affects organismal success. Within a species, natural variability produces individual mice that differ in aggression and flies that implement different foraging strategies (7, 8). Mutant screens have also uncovered genes associated with differences in locomotion, courtship routines, and sleep patterns, among other complex behaviors (2, 9–11).

Behavioral variability can be advantageous as a bet-hedging strategy against unstable environmental conditions(12). Phenotypic variability in behaviors ranging from birdsong to escape trajectories can increase individual success, but also fitness in a population, suggesting that variability itself can be a selectable trait. Experiments in *Drosophila melanogaster* demonstrate that the degree of behavioral variability in some locomotor behaviors is partially controlled by genetic expression of proteins such as *teneurin-α*, a cell adhesion molecule. Additionally, silencing a subset of neurons in the *Drosophila* central complex modifies the degree of variability of locomotor behavior (13). Differences in neurodevelopment and synaptic connectivity can also result in behavioral variability (14). Together, these experiments suggest that factors at both at the population (genetic) and individual (neuronal) levels contribute to behavioral variability.

Fruit flies live in dirty environments, from laboratory vials to rotting fruit, and perform grooming actions to remove accumulated particulate. This behavior has been observed in several drosophilid species and is important for social behavior and survival (15–17). Past work has demonstrated that the leg movements used in grooming are stereotyped, but the sequences of actions are flexible as opposed to fixed. While the rules, or syntax, underlying grooming do exhibit observable structure in flies (18, 19) and in mice (20), different sensory experiences and life histories may influence grooming behavior. These results lead us to ask: *how much variation and variability in fruit fly grooming is under genetic control*?

To address this question, N = 390 male flies were covered in dust and their grooming behavior was recorded for approximately 30 minutes each (Figure 1A). We analyzed flies from five drosophilid species (*melanogaster*, *santomea*, *sechellia*, *simulans*, and *erecta*), which are genetically distinct — separated by millions of years of evolution — and inhabit different ecological niches. We also examined four common *melanogaster* lab stocks (Canton-S, Oregon-R, Berlin-K, and w1118), and several isogenic lines derived from these parent stocks in our laboratory.

**Fig. 1.**
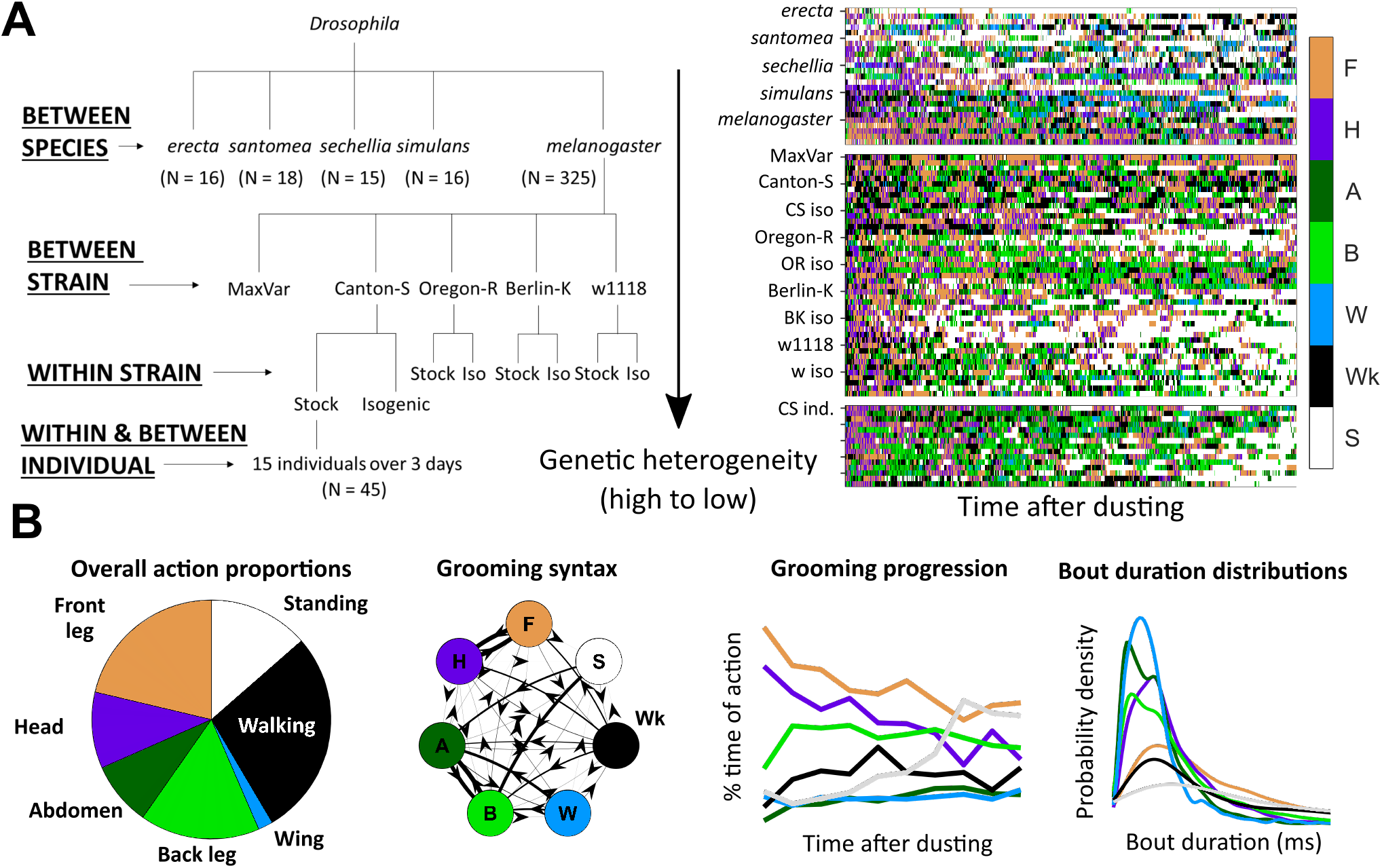
Grooming variability dataset and analysis overview. A. In total, *N* = 390 male flies were dusted and their grooming activity was recorded for approximately half an hour each. Five drosophilid species, four *melanogaster* stock lines, one interbred *melanogaster* line, and six isogenic *melanogaster* lines were analyzed for similarity, differences, and variability in grooming behavior. On the left is a schematic of the different drosophilid groups included in this analysis. Higher levels of the tree indicate higher levels of genetic diversity (scale is relative, not absolute). On the right is a sample of ethograms generated by automated annotation of grooming video. Color indicates the occurrence of the five grooming actions (F = front leg cleaning, H = head grooming, A = abdomen grooming, B = back leg cleaning, W = wing grooming), Wk = walking, and S = standing. B. Features scored from ethograms provide summary representations of grooming behavior. Shown on the left, the proportion of time spent in different actions, overall or over time, provides the coarsest description of the behavioral response to a dust stimulus. Regardless of genotype, all flies exhibit variable (not fixed) action sequences consisting of the same set of five grooming actions, walking, and standing after exposure to irritant. Next, grooming transition probabilities (syntax) describe the likelihood of performing consecutive actions. Arrow directions and thicknesses represent the probability of performing an action, given the identity of the previous action. Shown next is an example grooming progression, which depicts the proportion of time spent in each action over a sliding window. Most flies follow a typical grooming progression pattern: initial anterior grooming followed by increased posterior grooming. The amount and timing of walking and standing, however, can vary significantly between flies. Finally, action (bout) duration distributions describe the range of action lengths. All example features shown here are scored from Canton-S flies.

To analyze this large data set, we employed tools from computational ethology (21). An automated behavioral recognition system (ABRS, (22)) was used to classify fly behavior into one of five grooming actions (front leg cleaning, head grooming, abdomen grooming, back leg cleaning, wing grooming) and two non-grooming actions (walking and standing). As a note, head grooming consists of actions that use the front legs to clean the antennae, eyes, and face; sub-movements such as these were not easily detectable using the recording methodology employed here, so analysis was restricted to coarser spatiotemporal scales. After generating ethograms (behavioral time series records) for each fly, several grooming features were extracted (Figure 1B). We used classification analysis and various measures of stereotypy to quantify variation and variability of these characteristics.

We found that several features of grooming are conserved across different drosophilid species. Automated classifiers labeled actions with high accuracy for all species, indicating that the movement primitives that make up grooming are stereotyped. Flies tended to groom anterior body parts before posterior portions, showing a common progression. In all cases, grooming sequences were not strictly fixed, with no clearly identifiable long repeats of actions.

We evaluated different grooming features by comparing their average values between groups and their range within groups, using the concepts of *variation* and *variability* as defined by Ayroles et al. (11). Variation refers to differences in phenotype between genetically distinct populations, such as drosophilid species. For example, inter-species variation in drosophilid larval digging behavior has been identified based on the duration distributions of “dives” into an agarose substrate (23).

In contrast, variability describes how a trait differs within a genetically similar population. Intra-species variability has been characterized in the locomotor behavior of fruit flies, as some populations exhibit a wide range of locomotor behaviors, whereas others exhibit a narrower range (11).

Here, inter-species comparisons were used to look for variation in grooming features (such as time spent in different cleaning behaviors and grooming syntax) between flies with large genetic differences. Species differ in several ways, and the syntax differences are significant, as demonstrated by the fact that the transition probabilities among movements can be used to reliably classify individual flies by species.

Next, we looked for variation within *melanogaster* stock lines using similar analyses. Although grooming features were more similar between *melanogaster* stocks than between species, stock lines did exhibit detectable differences in grooming, largely related to overall activity levels (e.g., after dusting, some stocks spend a large proportion of time grooming, while others walk or stand more). Male and female flies also differed in activity levels, but to a lesser extent than between stocks.

Genetic differences among drosophilid species and strains may underlie variation in the syntax of their cleaning movements, but all flies show variability in the exact sequence of those movements. We hypothesized that genetic heterogeneity might contribute to the magnitude of this variability. By interbreeding or isogenizing *melanogaster* lab strains, we generated stocks with high and low genetic diversity, but we find that all groups exhibited similar variability in measured grooming features.

Furthermore, flies tested on sequential days revealed that the extent of within-fly differences in syntax were similar to between-fly differences; flies were no more similar to themselves over time than they were to other flies on a given day. Finally, flies stimulated using optogenetic manipulation to induce grooming exhibited increased stereotypy, but within-individual grooming variability between stimulation sessions was not fully abolished. These data show that genetic heterogeneity plays a limited role in the variability behavior, and that differences in sensory experience contribute but do not account for all observed variability in grooming behavior. The widespread nature of grooming variability suggests that it may be an important feature of the behavior, but our experiments indicate the need to search for alternative causes, perhaps including developmental stochasticity, differences in internal state, or noisy circuit dynamics.

## Results

### Drosophilids exhibit a robust grooming response but different syntax after irritant exposure

Across all *Drosophila* groups, five grooming actions were observed consistently, indicating a conserved behavioral response (Supplementary Figure 1). Previous work showed that these actions are sufficiently stereotyped to be reliably classified by manual and automated annotation in *melanogaster* (Automated Behavioral Recognition System ABRS, see Methods) (19, 22). Here, the ABRS classifier was validated on training data for each species, showing comparable accuracy (Methods). No novel species-specific grooming actions were detected. Although some fine-scale movement differences may occur among species, they are beyond the spatial and temporal resolution of the current video and unlikely to affect the analysis of transition probabilities presented here.

To quantify the behavioral response to dusting, the proportion of time spent grooming (as opposed to walking and standing) was calculated for each fly (Figure 2A). The grooming distributions between species were all statistically different (Wilcoxon rank-sum test, *p* < .05 after Bonferroni correction), but all species spent at least 35% on average. In this analysis, a single stock line (Canton-S, N = 18) was used as the representative *melanogaster* group. Full action proportion distributions are shown in Supplementary Figure 2.

**Fig. 2.**
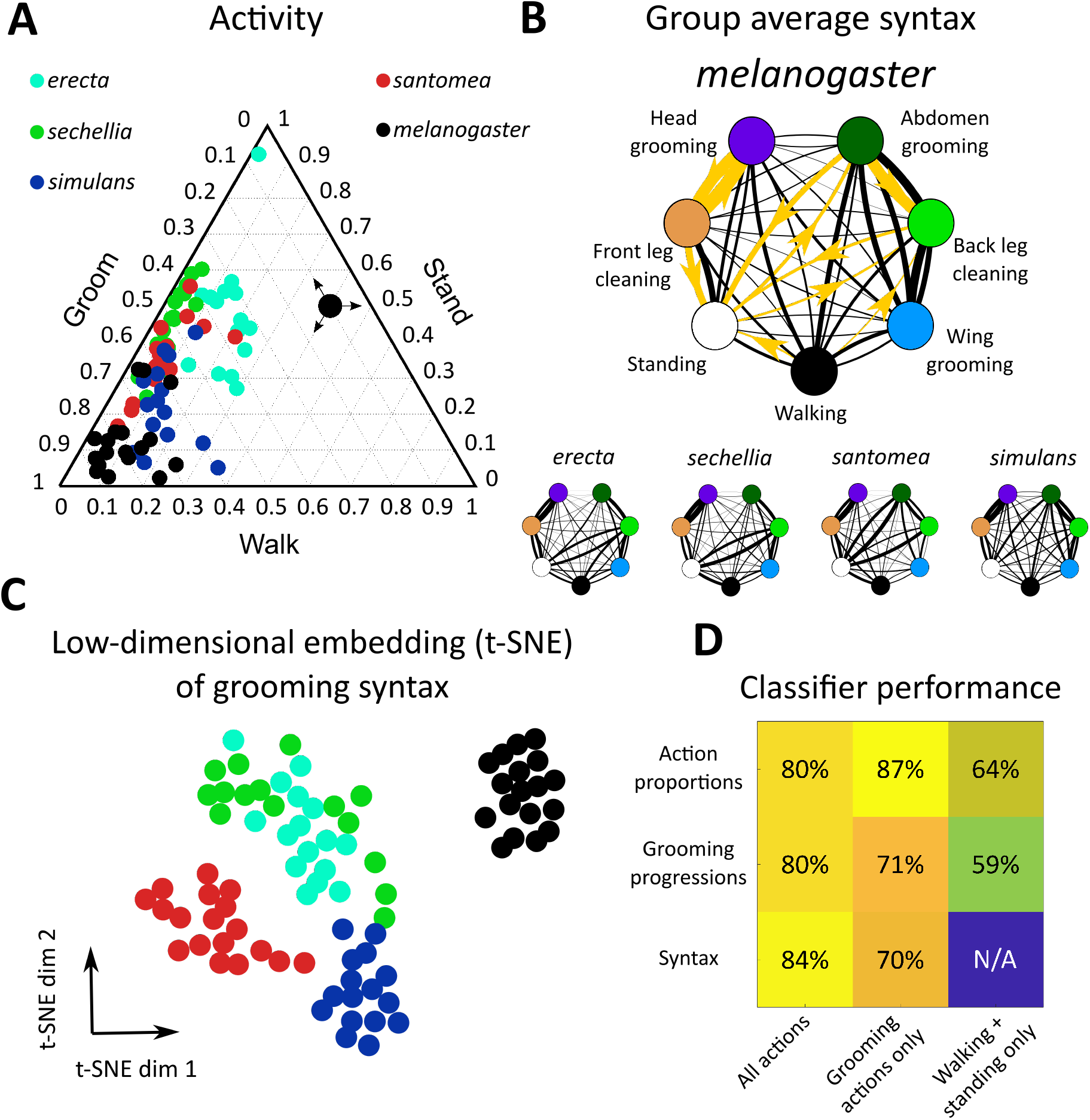
*Drosophila* species share grooming features but exhibit between-species variation in action proportions and syntax. A. Dusting elicits a conserved grooming response across drosophilids. Shown is a ternary plot of action proportions for each species examined here. Colored points represent a single fly, with color indicating species. The large black point with arrows indicates how to read activity proportions; the example point corresponds to 10% grooming, 40% walking, and 50% standing. B. Drosophilid species produce a probabilistic grooming sequence (as shown in Figure 1A), which can be characterized by the transition probabilities (syntax) between grooming actions (as represented in Figure 1B, calculated as in as in (19)). The mean syntax for each species is depicted as a graph, with nodes representing grooming actions and edges indicating transition probability. Thicker edges indicate higher probabilities. On the *melanogaster* syntax graph, the 10 action transitions exhibiting the largest magnitude differences between *melanogaster* and non-*melanogaster* species are highlighted in gold. These differences are identifiable in anterior motif transitions, which use the front legs to perform grooming actions. Species also differ in their transitions between posterior grooming actions and non-grooming actions (walking and standing) C. Each fly’s 42-dimensional syntax vector was plotted in two dimensions after dimensionality reduction using t-SNE. t-SNE preserves local distance structure, indicating that tightly grouped clusters of points are similar to one another. In this case, dimensionality reduction reveals that drosophilid species exhibit significant differences in grooming syntax, as syntax vectors congregate by color. D. Classification analysis confirms the qualitative clustering observed in C. Shown is a heat map of accuracy rates of 5-possibility multinomial logistic regression classifiers trained on grooming features. For these samples, classification at chance would be 20%. Consistent classification accuracy values >20% indicate that species are highly separable by grooming features. Simple grooming features, such as behavioral proportions and progressions, classify individuals by species with high accuracy when grooming actions are included. Classification using only non-grooming actions (walking and standing) still yields classification above chance, indicating that species differ significantly in their overall activity levels. Syntax also allows for accurate classification, particularly when all action transitions are considered.

Grooming was then examined in more detail by considering all seven actions and the progression of those actions over time, as in Figure 1B. These species exhibited a qualitatively similar grooming progression characterized by initial anterior grooming followed by increased posterior grooming, walking, and standing, but the relative proportions and timing of these behaviors over time differed (Supplementary Figure 3).

Grooming syntax (the transition probabilities between discrete behaviors) was calculated from the ethogram of each dusted fly (*N* = 390). With seven behavioral states, 42 transitions were possible, excluding self-transitions. Thus, syntax was represented as a 42-dimensional vector for subsequent classification analysis and visualization. Grooming syntax across all flies exhibited high transition probabilities within the anterior grooming motif (front leg cleaning, head grooming) and posterior grooming motif (abdomen grooming, back leg cleaning, wing grooming). The average syntax for each species is illustrated as a weighted, directed graph in Figure 2.

Finally, continuous action duration distributions (e.g., the distribution of how long each head grooming action was) were calculated from ethograms. Distributions of grooming action durations were similar across species and had probability peaks between 500 and 750 ms (Supplementary Figure 4).

Several significant differences in grooming features between *melanogaster* and non-*melanogaster* species were identified. Supplementary Figure 5 illustrates differences in overall action proportions, around 36% of which differed between species. To compare syntax, transition probability distributions for each action transition (e.g., head cleaning to front leg rubbing) were compared between species in a pairwise manner. 38 of 42 unique syntax elements (90.5%) were significantly different between at least two species (Wilcoxon rank-sum test, p<0.05 after Bonferroni correction). Overall, 119 of 420 (28%) of pairwise syntax comparisons revealed differences between species.

Of these syntactic differences, 71 (60%) occurred between *melanogaster* and non-*melanogaster* species. In particular, posterior motif grooming transitions (transitions between abdomen grooming, back leg rubbing, and wing grooming) were consistently significantly different, on average, as were transitions between back leg rubbing, standing, and walking. Figure 2B illustrates these syntactic differences.

Figure 2C depicts a low-dimensional embedding of species syntax using t-SNE. This visualization suggests that different species possess distinguishable syntax, as points are aggregated by species. Low-dimensional visualizations of all grooming features are illustrated in Supplementary Figures 5 and 6.

Classification analysis was applied to grooming features to verify this interpretation and quantify the degree of variation between species. Multinomial logistic regression classified flies by species according to behavioral proportions, progressions, and syntax with >80% accuracy (Figure 2D). Notably, classification was also possible with accuracy significantly above chance when only considering the proportions and progressions of non-grooming actions, walking and standing, indicating that species also vary in their overall activity levels.

Finally, entropy rates were calculated from syntax transition probabilities to quantify the degree of stereotypy in behavior. An entropy rate of zero would indicate complete stereotypy and perfectly predictable, repeated action sequences, while in this calculation, an entropy rate of one indicates an approximate 37% probability of correctly predicting the next action in a sequence (see Methods). Supplementary Figure 7 shows that all species possess average entropy rates between zero and one, demonstrating that grooming sequences are neither fixed nor truly random. *melanogaster* flies possessed the lowest entropy (highest degree of stereotypy) due to high transition probabilities between head cleaning and front leg rubbing (Figure 2B). In summary, drosophilid species exhibit variation in grooming behavior detectable across several different grooming features.

### *melanogaster* strains exhibit variation in grooming behavior

Next, standard *Drosophila melanogaster* lab strains (Canton-S, Berlin-K, Oregon-R, w1118) were analyzed for differences in grooming features (full ethograms shown in Supplementary Figure 8). Behavioral proportions, progressions, and syntax differed between stocks, allowing for classification moderately above chance levels. Comparisons of grooming features can be found in Supplementary Figures 9 to 12.

Overall the proportion of time grooming could account for most of the differences observed between stocks. Figure 3A shows a ternary plot of activity, showing that Canton-S flies spend more time walking than other stocks. A t-SNE embedding of the grooming syntax of *melanogaster* stocks is depicted in Figure 3A. Similar to the species analysis, all action transition probability distributions were compared in a pairwise manner to look for variation in syntax (examples in Figure 2A).

**Fig. 3.**
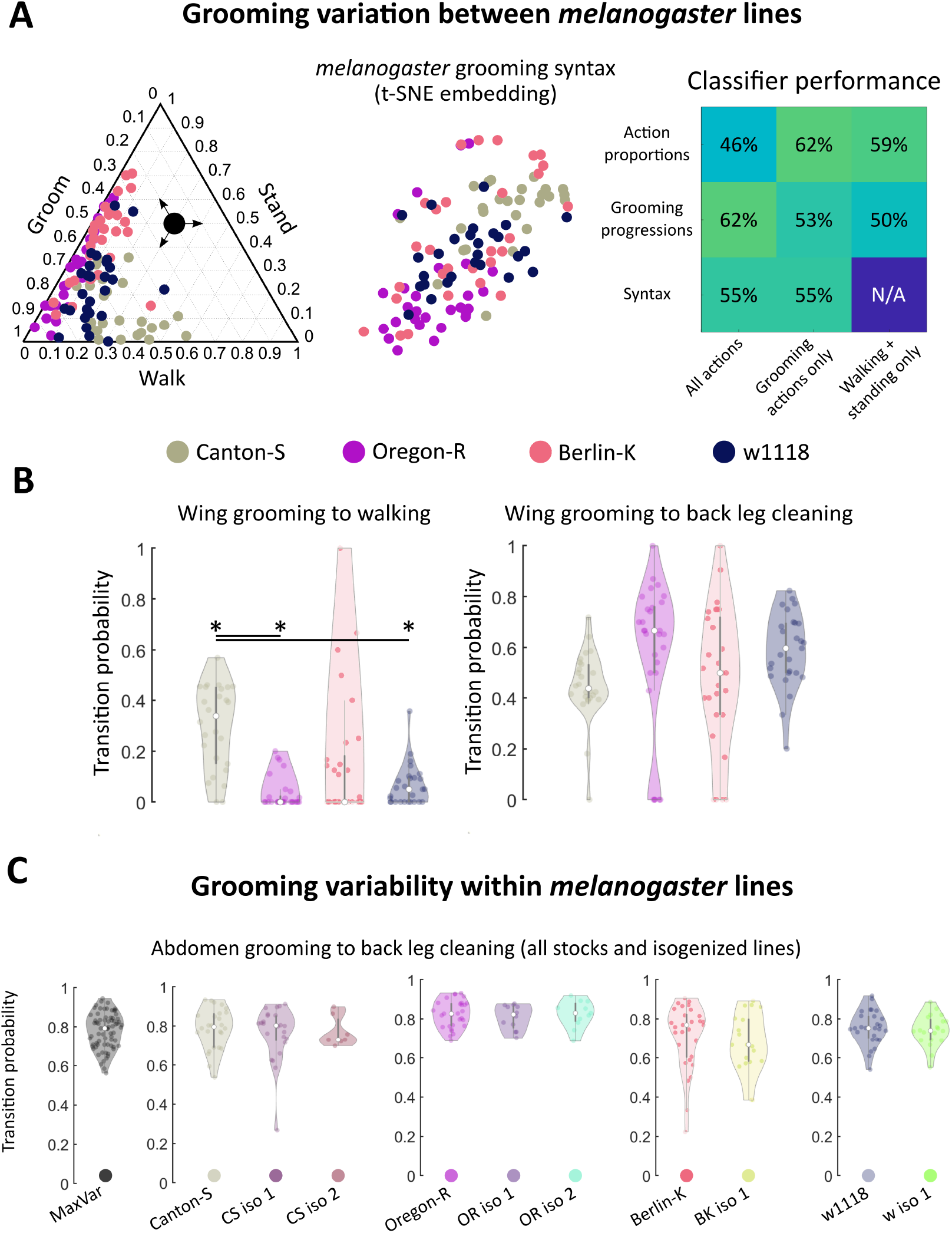
Within *melanogaster*, different stocks differences in syntax activity levels. Genetic homogeneity does not correspond to behavioral stereotypy. A. *melanogaster* stocks exhibited variation in grooming syntax, though many features were shared. On the left is a ternary plot of grooming, walking, and standing proportions for each stock, similar to Figure 2A. Colored points represent individual flies. Shown in the middle is a t-SNE plot of grooming syntax vectors, as in Figure 2C. The high degree of overlap in both of these plots illustrates that grooming responses are qualitatively similar between some individuals of different stock lines. Classifier performance (similar to that shown in Figure 2D) is shown on the right. For these data, classification at chance is 25%. Performance above chance is still possible for stock lines. Classification performs similarly well for grooming features regardless of their complexity; using just walking and standing behavioral proportions provides similar discriminability as using the full syntax. B. Most grooming syntax elements were similar between *melanogaster* stocks, but Canton-S flies walked more than other stocks. Due to differences in activity levels, some walking-related syntax elements differed between Canton-S flies and other stocks. Of the significantly different transitions, only two were within-motif transitions while the rest consisted mostly of transitions to and from walking and standing (Supplementary Figure 12). Shown on the left are the wing grooming to walking transition probability distributions for each *melanogaster* stock line. Significant differences in these distributions were observed between lines. On the right, distributions for a posterior grooming transition are shown; the vast majority of grooming action transition distributions did not differ due to their large variances. C. Variances of action transition distributions for stock lines, lines bred for maximum genetic heterogeneity (MaxVar), and lines bred to minimize genetic heterogeneity (iso) were compared. Genetic homogeneity did not correspond to behavioral variability. Shown as an example are the distributions of abdomen grooming to back leg cleaning transitions. MaxVar flies did not exhibit a higher degree of variability (as measured by the variance of transition distributions) than stock lines. Isogenized lines did not exhibit a lower degree of variability than their parent stocks.

Only 19 of 42 unique syntax element comparisons (45%) differed significantly between any two stocks and, of these, only two within-motif transition (both posterior motif) differed significantly. Within-motif syntax elements are of particular interest because they represent the most common, most highly stereotyped action transitions observed across flies of all genotypes (see Figure 2A and B for visualizations of these transitions). The syntax element exhibiting the greatest statistically significant difference was the wing grooming to walking transition, shown in the middle of Figure 3B. The other significantly different transitions also mostly involved transitions to and from walking and standing (Supplementary Figure 12).

Classification accuracy was moderate but above chance for all features examined; as expected, variation within *melanogaster* was less pronounced than variation between species (compare Figures 2D and 3A). Variation within *melanogaster* stocks appears to be due to differences in overall activity levels, as classification using only non-grooming features (walking and standing proportions and progressions) yielded results similar to classification using full grooming behavior syntax. This is illustrated by the fact that Canton-S flies’ higher propensity to walk after grooming their wings is reflected both in their syntax and grooming proportions in Figure 3.

Within Canton-S, activity levels separated male and female flies, as male flies tended to walk more than females (Supplementary Figure 13). Male and female flies also possessed somewhat different syntax; classification by syntax was 71%, where chance levels would be 50% for this comparison. This level of accuracy is higher than what was achievable when classifying *melanogaster* stock lines using syntax, but lower than the same comparison for interspecies data.

Since all flies examined showed variability in syntax, we wondered whether the extent of this variability differed among species or strains. Figure 3B also illustrates the high degree of variability in *melanogaster* syntax. The wing grooming to back leg cleaning transition exhibited the largest difference between median values of any syntax element (comparison of Canton-S and Oregon-R yielded this difference), but none of these distributions possessed detectable statistical differences due to their concomitantly large variances.

### Grooming behavioral variability is similar across *melanogaster* genotypes

To examine the relationship between genetic heterogeneity and behavioral variability, each *melanogaster* lab strain was compared to lines bred to maximize genetic heterogeneity (MaxVar) or minimize genetic heterogeneity (isogenic lines). If variability in grooming syntax within a population is strongly related to genetic heterogeneity, we would expect populations with large genetic heterogeneity to also contain flies with widely varying syntax structures.

All lines, regardless of genetic heterogeneity, exhibit variable grooming (Supplementary Figure 14). To quantify variability, the variances of grooming transition probability distributions were calculated and compared. Only 6/252 (2.4%) transition probability distributions possessed statistically significantly different variances between MaxVar, Canton-S, and the isogenic lines out of all possible pairwise comparisons (Levene’s test, *p* < .05 after Bonferroni correction). Moreover, none of these differences corresponded to within-motif transitions, indicating that variability of common transitions is similar regardless of genetic heterogeneity in a population. These findings also held for Oregon-R (8/252), Berlin-K (18/108), and w1118 (2/108) stock and isogenic comparisons.

Figure 3C provides the transition probability distributions for the most common posterior motif transition (abdomen grooming to back leg cleaning) for all stocks and stock-derived isogenic lines. This transition exhibits wide variability in many populations and even populations with smaller variability (CS iso 2) are not different enough to achieve statistical significance after accounting for multiple hypothesis testing.

We also examined stock lines derived from selected wild isolates (24) to determine if these showed more or less grooming variability, as measured by syntax element variance values and Markov entropy. The variability within wild isolate lines is comparable to that within lab stocks (Supplementary Figure 15).

Finally, we analyzed 15 Canton-S flies that were recorded over three consecutive days after irritant exposure. Since a given individual’s genome remains constant through the three trials, we could isolate the magnitude of grooming variability that is due to differences in sensory experience (since the dusting protocol does not allow for perfect replication of sensory experience) and life history (since flies will have been exposed to the same irritant several times by the end of the experiment). Ethograms from three example flies are provided in Figure 4A (full ethograms are shown in Supplementary Figure 16).

**Fig. 4.**
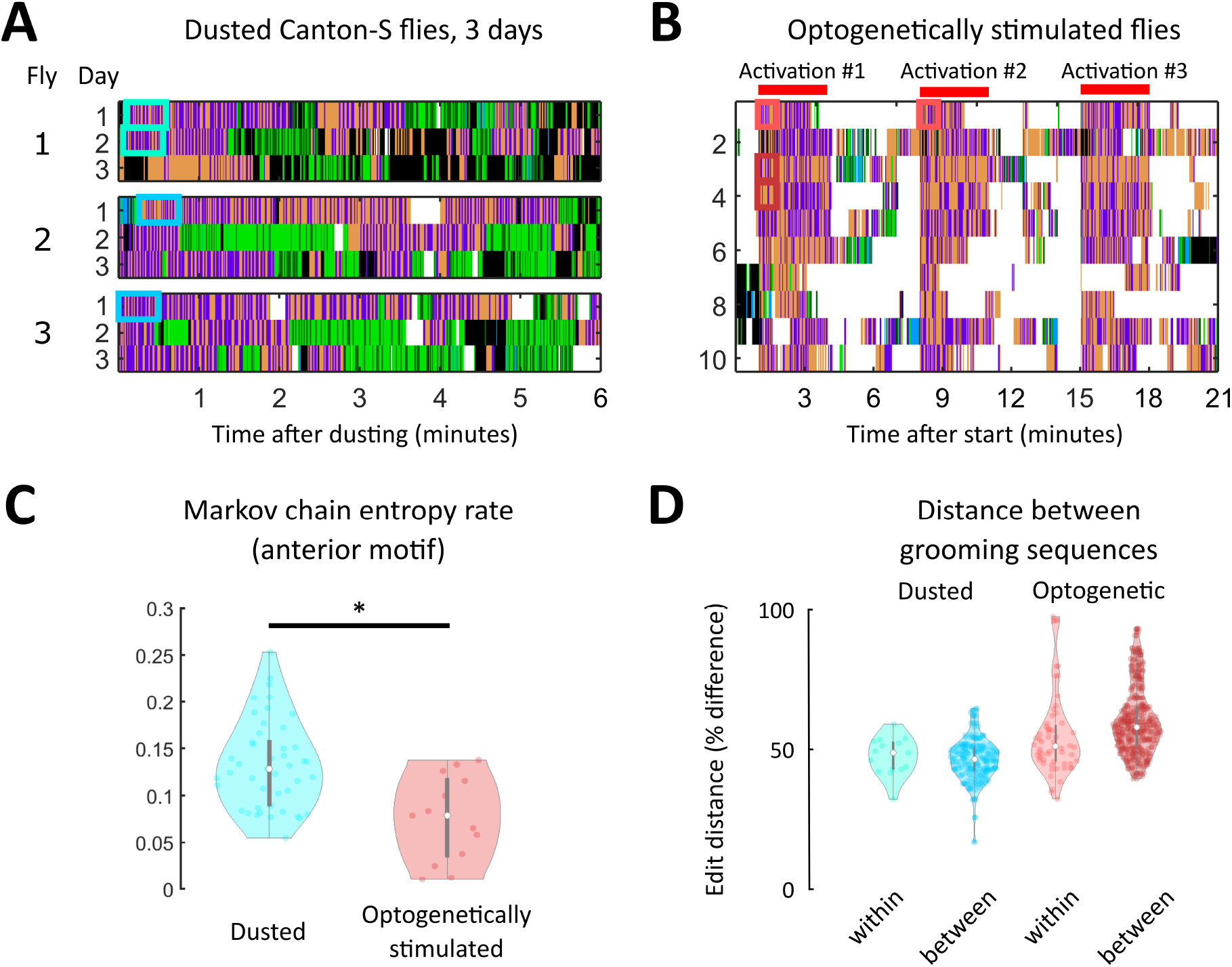
Within-individual grooming differences suggest that non-genetic factors account for a significant portion of variability in behavior. A. Portions of ethograms from three Canton-S flies observed on consecutive days after dust irritant exposure. The differences in ethograms on consecutive days indicate that non-genetic factors must account for some amount of grooming variability. B. Shown are ethograms of 10 Bristle-spGAL4-1 > CsChrimson flies (25). Flies were optogenetically stimulated to induce anterior grooming in three separate three-minute windows, indicated by the red bars. Between these windows, flies still exhibit within-individual grooming variability even though the sensory experience is more uniform than repeated dust exposure. C. Markov chain entropy, a measure of grooming stereotypy, was calculated from anterior grooming syntax. Optogenetically stimulated flies (right) exhibited lower entropies, corresponding to a higher degree of stereotypy, than dusted flies (left). However, optogenetically stimulated flies still exhibited differences in stereotypy between stimulation windows, implicating sources of grooming variability beyond genetic and sensory influences (Supplementary Figure 19). D. To assess grooming stereotypy, edit distance between anterior motif repeats was computed. For dusted within-fly comparisons, we computed the edit distance between the first continuous anterior motif sequence lasting thirty seconds on consecutive days (light blue). For between-fly comparisons, we computed the edit distance between the first continuous anterior motif sequence lasting thirty seconds on the first day of experiments (dark blue). For all optogentically-stimulated flies, we computed two similar comparisons: within-session (i.e., comparing the sequences labeled “Activation #1” and “Activation #2” in panel B; light red) and between-fly (i.e., “Activation #1” for each fly; dark red). For each comparison listed, the median edit distance computed corresponded to around 50% of the sequence length, demonstrating the low degree of stereotypy present in grooming sequences.

Flies exhibited some longitudinal grooming trends, as the total amount of grooming decreased between the first and third days of the experiment. However, the time to completion of 50% of their total grooming did not decrease, suggesting that flies are not simply grooming quicker, but rather are grooming less consistently over time (i.e., punctuating grooming bouts with more walking and standing) (Supplementary Figure 17). Importantly, intra-individual variability in syntax across three sessions was of the same magnitude as inter-individual variation in syntax (Supplementary Figures 18 and 19); that is, flies were no more similar to themselves over time than they were to other flies on a given day. This suggests that non-genetic factors account for a significant proportion of grooming variability.

### Standardizing sensory experience does not abolish grooming behavioral variability

To probe the sensory contribution to within-individual variability, we used optogenetic stimulation to induce anterior grooming. 20 Bristle-spGAL4-1 > UAS-CsChrimson flies were tested (25). Figure 4B provides ethograms from this experiment, with red bars indicating the three stimulation windows. Even when sensory experience was controlled in this way, flies exhibited variability in their grooming response.

Grooming stereotypy was again quantified using the entropy rate of the grooming syntax. The entropy rate for optogenetically stimulated flies was lower than for dusted flies (*p* < .05, Wilcoxon rank-sum test), indicating a higher degree of stereotypy in grooming (Figure 4C). We nonetheless observed within-individual variability between stimulation windows, indicating that standardization of sensory input does not fully abolish grooming variability. Supplementary Figure 20 quantifies differences in entropy between sessions for three example flies. In addition, optogenetic stimulation resulted in strong anterior motif grooming behavior, rendering all flies’ transition probabilities very similar (Supplementary Figure 21).

Finally, grooming stereotypy was characterized using edit distance between anterior motif repeats. This metric, used commonly in bioinformatics, describes the difference between two DNA sequences by calculating the minimum number of base pair substitutions, additions, or deletions that would be necessary for the sequences to be identical. Identical sequences would have an edit distance of zero between them, while maximally different sequences would have an edit distance equivalent to the total sequence length (see Supplementary Methods for details). For all dusted flies, we calculated the edit distance within flies across consecutive days to assess whether flies possess stereotyped repeats. For these comparisons, we compared the first continuous anterior motif sequence lasting at least thirty seconds on consecutive days. This particular comparison was chosen to standardize the amount of dust present on the fly to the greatest extent possible and the short-term grooming history for each sequence, and to ensure that each sequence was long enough to exhibit stereotypy if it exists. Anterior motifs were chosen because they consist of only two actions with high transition probabilities between them, making these sequences the most likely candidates for exhibiting stereotypy. These comparisons yielded a minimum edit distance corresponding to a 39.6% difference between sequences. A similar calculation was made between flies, using the first continuous anterior motif sequence lasting at least thirty seconds on the first day of experiments. These comparisons yielded a minimum edit distance corresponding to a 41.6% difference between sequences.

Similar calculations were then performed for all optogenetically-stimulated flies. Within-fly comparisons (i.e., comparing the sequences labeled “Activation #1” and “Activation #2” in panel B) yielded a a minimum edit distance corresponding to a 31.5% difference in sequences. Between-fly comparisons (i.e., “Activation #1” for each fly) yielded a a minimum edit distance corresponding to a 42.8% difference in sequences. Together, the low degree of stereotypy present in grooming sequences within and between both dusted and optogenetically stimulated flies shows that grooming sequence variability is present even when genetics, sensory input, and behavioral history are controlled to the greatest extent possible within this experimental paradigm.

## Discussion

Here, we analyzed fly grooming behavior in five different drosophilid species and four common *melanogaster* stocks to investigate the relationship between genetic heterogeneity, variation, and variability. Large genetic differences (species-level) correspond to identifiable differences in several grooming features, including the rules governing action transitions known as syntax. Within *melanogaster*, stock lines also exhibited smaller variation in grooming syntax, as well as differences in overall activity levels. All flies showed variability in the details of the grooming movement sequence, but increased genetic heterogeneity did not correspond to increased behavioral variability. Analysis of 15 Canton-S flies recorded over consecutive days showed that intra-individual and inter-individual comparisons had similar — high — levels of variability. Optogenetically stimulated flies also exhibited intra-individual variability in grooming behavior, but less. Taken together, these results demonstrate that large genetic differences result in distinguishable grooming phenotypes (variation), but that genetic heterogeneity within a population does not necessarily correspond to an increase in the range of grooming behavior (variability).

### Genetic influences on behavioral variation and variability

Advantageous behavioral phenotypes that are under genetic control can be selected over evolution to produce populations with differing behaviors (variation). Here, we identified significant inter-species variation in grooming syntax, suggesting a genetic basis for group differences in grooming behavior. Some species differ from *melanogaster* in their propensity to perform anterior grooming actions; the anterior motif actions are significantly less strongly coupled in non-*melanogaster* drosophilids, suggesting that anterior neuronal circuitry and sensory physiology may differ. We also identified differences in grooming behavior between commonly used *melanogaster* stock lines and between male and female Canton-S flies; most of these differences relate to overall activity levels.

Variability itself is a trait that can also be selected for. At the individual level, randomizing escape trajectories can be beneficial for escaping predators (26), and diversity in search paths can be useful when a group is foraging for food. The fate of the passenger pigeons, hunted to death while flocking together, illustrates the dangers of behavioral homogeneity (27). The degree of variability in behavior can be selected for as a bet-hedging strategy against unstable environmental conditions (12, 28). Genetic factors contribute to variability in fly visual, olfactory, and locomotor behaviors (11, 29, 30).

The prevalence of variability in *Drosophila* grooming sequences suggests that non-stereotyped grooming may be helpful in removing diverse distributions of debris. We examined whether greater genetic heterogeneity within a population corresponded to greater behavioral variability but did not detect any significant impact.

A recent investigation of unstimulated behaviors in different *Drosophila* species detected differences in spontaneous grooming between species and among individuals within a species (31). Using similar methods, they accurately assigned individuals into species categories and assessed variability among individuals. While our studies focused on different aspects of genetic contributions to behavior, our findings are complementary: drosophilid species show differences in stimulated grooming behaviors as well, suggesting genetic control, but individuals within a species show variability in grooming, indicating that factors other than genes can influence aspects of the behavioral sequence. Hernandez et al. propose that over the long timescales measured in their assay, internal states may explain the observed fluctuation in action transition probabilities. In the shorter timescales we assayed, where flies are responding acutely to dust, we attribute the variability to inherent flexibility in the behavior itself, produced by differences in sensory input and/or intrinsic stochasticity in the neurons or circuits that coordinate the action sequences. These views are not in conflict. Together, we establish that variability in grooming is widespread — potentially even advantageous — and that both genetic and non-genetic factors influence its expression.

Variability also encompasses individuality in animal behavior, typically defined as a trait-like feature that persists stably over several observations. Individuality has been identified in fruit fly turning decisions (32), mouse roaming behavior (33), and bumblebee foraging (34), among others (30, 35). In both dust-induced and optogenetically initiated grooming, we did not find evidence for individuality in grooming sequence action patterns at the resolution we analyzed, but this may be because small contributions from individual tendencies are outweighed by the large amount of variability in the behavior as a whole arising from other causes.

### Environmental and stochastic influences of behavioral variation and variability

Our analysis of genetic contributions to behavioral variation and variability in the grooming suggests that at the species level, flies show significant differences in the grooming sequence, especially in the syntax of transition probabilities, that allow accurate classification. Differences in grooming behavior between common lab wild-type stocks also support classification, but the accuracy is lower and the effect size of the differences is smaller.

Genetic factors have been implicated in spontaneous (i.e., unstimulated) grooming behavior in *Drosophila melanogaster* (36) and in other drosophilid species (31). Our results demonstrate that this is true for dust-induced grooming as well. Both spontaneous grooming (31) and dust induced grooming show individual-to-individual variability within a species. The prevalence of sequence flexibility in all species and in controlled experimental conditions suggests that variability itself is a feature of grooming behavior, not a bug. Individuals with overly rigid grooming sequences might not respond as effectively to changing environmental conditions, such as different kinds of debris or the presence of a potential mate or predator.

The precise genetic contributions to this property are still under investigation. Differences in developmental processes such as neural wiring or synaptic connectivity may contribute to behavioral differences between flies, but our experiments show that even individual flies exhibit variability in grooming over repeated trials with dust or optogenetic stimulation. This suggests that non-genetic factors such as sensory stimuli, internal state, previous experience, and circuit noise contribute to the variability we observe in grooming action sequences. The reduction of variability when sensory inputs are optogenetically controlled supports diversity of sensory stimulation as a contributor. The persistence of variability within individuals suggests that intrinsic stochasticity or noise within the neurons or circuits themselves may also play a role, which are possibilities which should be explored further.

## Materials and methods

### Genetic stocks

Canton-S, Oregon-R, Berlin-K, w1118, Bristle-spGAL4-1 (R38B08-AD; R81E10-DBD), and 20XUAS-CsChrimson-mVenus (attp18) stocks were obtained from the Bloomington Stock Center. Isogenic (more accurately, reduced genetic variability) stocks were made by crossing single males to double-balanced stocks and then back-crossing males to the double balancer stock to isolate single second and third chromosomes. Single pairs were mated to reduce variability of X and IV. ~ 2 independent isogenic lines from each *melanogaster* stock were generated; note that many attempts to isogenize result in lethality, as anecdotally reported by colleagues. Maximum Variability stocks were obtained by crossing each *melanogaster* strain to double balancers and then crossing the progeny together and selecting against the balancers. This allowed combination of chromosomes for all four strains. The progeny were allowed to interbreed for several generations to enable recombination in the females. Drosophilid species stocks were obtained from Tom Turner, UCSB, and the National Drosophila Species Stock Center (https://www.drosophilaspecies.com/).

### Data collection and processing

Grooming was induced and analyzed as described in Zhang et al. 2020 and Seeds et al. 2014 (18, 25). Three chambers were used in fly dusting assay: dusting chamber (24 well Corning tissue culture plate #3524), transfer chamber and recording chamber. Recording chambers were coated with Insect-a-slip (BioQuip Products Cat#2871A) to discourage wall-climbing and cleaned daily. Dust-induced grooming assays were performed in 21-23°C. 4-7 day old male flies were anesthetized on ice and transferred to the middle four wells of the transfer chamber. Flies were left in the transfer chamber for 15 minutes to recover. Approximately 5 mg Reactive Yellow 86 dust (Organic Dyestuffs Corporation CAS 61951-86-8) was added into each of the 4 middle wells of dusting chamber. For fly dusting, the transfer chamber was aligned with the dusting chamber. Flies were tapped into the dusting chamber and shaken 10 times. After dusting, flies and dust were transferred back into the transfer chamber. Transfer chamber was tapped against an empty pipette tip box to remove extra dust. Dusted flies were then immediately tapped into recording chamber for video recording. The entire dusting process was performed in a WS-6 downflow hood. Approximately 10 individuals were recorded for each genotype. 30 Hz videos were recorded for 50,000 frames (27.78 min) with a DALSA Falcon2 color 4M camera. A white LED ring right was used for illumination.

Optogenetic stimulation protocol is replicated from (25). Further details can be found in the Supplementary Methods. For each set of experimental comparisons (between species, within species, within individual), a single experimenter performed all dusting assays to eliminate experimenter-related differences that may arise.

Videos were processed through the Automated Behavior Recognition System (ABRS, (22)), trained on a classifier using *melanogaster* flies to generate ethograms. Grooming actions were described previously (37). Sub-movements of the grooming actions used in this analysis have not yet been rigorously described and may occur on time scales faster than the 30 Hz recording setup can reliably capture, so they were not considered in this work.

ABRS was used to generate ethograms. Briefly, the raw video frames were pre-processed to generate 3-channel spatial-temporal images (ST images). Features were extracted in three timescales and saved into different channels of ST images: 1. raw frame; 2. difference between two frames; 3. spectral features extracted from a 0.5 sec window. A convolutional network trained by ST images under different light conditions was then used to label the behavior identified in each frame. A different network was trained for classification of each species due to differences in body size and light conditions. All networks achieved > 70% validation accuracy within the training protocol, which reserved 20% of frames as test data after training. The first three minutes (5400 frames) of one ethogram per species were also manually annotated and compared to ABRS output. Manual annotation did not identify grooming actions that were unique to a subset of *Drosophila* species. This manual validation also achieved >70% accuracy for all species, which is similar to the agreement between human-annotated ethograms (22).

Finally, ethograms were denoised to only include grooming actions that persisted for longer than the approximate duration of one complete leg sweep. Here, we used a cutoff of 150 ms, and eliminated any actions shorter than this duration (fewer than 1% of bouts were removed under this criterion).

### Data analysis

All ethogram features were extracted using custom-written code in MATLAB 2019a. Grooming progression vectors were generated for each fly by calculating the proportion of each action in 10 non-overlapping windows (2.78 minutes each), yielding a 70-dimensional vector for each fly (10 windows with 7 behavioral proportions). Grooming syntax was defined as the first-order transition probabilities between actions. Syntax for each fly was calculated as described in (19). Bout duration distributions were generated as described in (19), using a normalized histogram with 20 bins of equal width for each behavior. Bin width was determined independently for grooming and non-grooming actions, as standing and walking exhibit longer tailed distributions than grooming actions. Thus, duration distribution vectors were 140-dimensional for each fly. Examples of progression, syntax, and duration distribution vectors can be found in the Supplementary Information.

Statistics for comparisons between grooming features were calculated using built-in MATLAB functions. t-SNE, and multinomial logistic regression classification analysis were performed using built-in MATLAB functions (Supplementary Information).

## Author Contributions

Experimental design: J.H.S. Data collection: J.H.S., N.Z.; data analysis: J.M.M.; manuscript writing: J.M.M.; manuscript editing: J.M.M., J.H.S., J.M.C.

## Acknowledgments

Stocks obtained from the Bloomington Drosophila Stock Center (NIH P40OD018537) were used in this study. Thanks to Karen Hibbard, Barry Ganetzky, and Claire McKellar for consultations on isogenization strategies. Thanks to Jordan Zawaydeh for data collection of individual fly data. Thanks to Kristin Branson, Matthieu Louis, Roian Egnor, and Primoz Ravbar for advice and comments on the manuscript.

