## Supplementary Information for "Variation and variability in *Drosophila* grooming behavior"

**CONTENTS**


Supplementary Methods 3

Classification analysis 3

Visualization 3

Markov chain entropy rate 4

Edit distance 4

Optogenetic stimulation 5

References 5

Supplementary Figures 6

### SUPPLEMENTARY METHODS

#### Classification analysis

To determine whether flies were distinguishable on the basis of several grooming features, logistic regression classification was applied following [1]. Specifically, nominal multinomial logistic regression classifiers were trained and tested using the MATLAB functions `mnrfit` and `mnrval` with a logit link function. Models were fit using leave-one-out cross-validation to prevent overfitting.

Separate classifiers were trained on each grooming feature (syntax, progression, and behavioral proportions) for comparisons within the following multi-class datasets: *erecta*, *santomea*, *sechellia*, *simulans*, and Canton-S; Canton-S, Berlin-K, Oregon-R, and w1118; Chance classification accuracies are given in the main text. Accuracy was percentage of labels predicted by classifiers that match the true class labels.

#### Visualization

To visualize high-dimensional grooming features in low-dimensional space, t-Distributed Stochastic Neighbor Embedding (tSNE) was used [2]. Briefly, tSNE performs dimensionality reduction by converting high-dimensional pairwise Euclidean distances into conditional probabilities (or similarities). These probabilities are given by

$$p_{j|i} = \frac{e^{-\|x_i - x_j\|^2/2\sigma_i^2}}{\sum_{k\neq i} e^{-\|x_i - x_k\|^2/2\sigma_i^2}} \tag{S1}$$

where $\sigma_i$ is the variance of a Gaussian distribution centered at $x_i$. These probabilities represent the likelihood that any two points would be nearest neighbors in low-dimensional space, assuming a Student-t distribution of pairwise distances between points. The low-dimensional counterpart joint probabilities, then, are given by

$$q_{ij} = \frac{(1 + \|y_i - y_j\|^2)^{-1}}{\sum_{k\neq l}(1 + \|y_k - y_l\|^2)^{-1}} \tag{S2}$$

where $y_i$ is the low-dimensional embedding of $x_i$. To create a map between high and low dimensions, the Kullback-Leibler divergence between P, the high-dimensional joint probability distribution, and Q, the low-dimensional distribution, are minimized using the cost function

$$\sum_i KL(P_i \| Q_i) = \sum_i \sum_j p_{j|i} log \frac{p_{j|i}}{q_{j_i}} \tag{S3}$$

where $P_i$ is the high-dimensional conditional probability distribution of the point $x_i$ and $Q_i$ is its low-dimensional counterpart. A non-deterministic solution to this mapping cost function is found using gradient descent. Here, the `tsne` function in MATLAB was used to perform this computation. 5000 gradient descent steps were used, as solutions were found to converge sufficiently using that number of steps. The perplexity value, a hyperparameter that approximates the effective number of neighbors of a point, was set between 15 and 30. Note that typical perplexity values range between 5 and 50.

**Markov chain entropy rate**

Using first-order transition dynamics to define syntax, the Markov chain entropy rate was calculated as a measure of stereotypy in grooming. An entropy rate of 0 indicates that grooming transitions are perfectly predictable, while higher values indicate decreased stereotypy. Entropy was defined as

$$H = -\sum_{ij} \mu_i P_{ij} ln(P_{ij}) \tag{S4}$$

where $P_{ij}$ is the transition probability from action $i$ to action $j$ and $\mu$ is the steady-state action distribution.

**Edit distance**

To assess grooming stereotypy, 30-second long (900 frames) anterior motif sequences were converted to text strings. Edit distance was then calculated in a pairwise manner from these strings using the MATLAB function `editDistance`. This function returns the lowest number of insertions, deletions, and substitutions required to render the strings identical.

**Optogenetic stimulation**

Protocols for optogenetic stimulation of grooming are reproduced here in brief from Zhang et al. (2020).

After cold anesthesia, flies were left to recover in recording a chamber for at least 20 min. Custom-made LED panels (LXM2-PD01-0050, 625nm) were used for light activation from below. 20 Hz 20% light duty cycle was used in all experiments. Light intensity was measured by Thorlabs S130VC power sensor coupled with PM100D console. The light intensity used for Bristle-spGAL4-1 flies was 0.84 $mW/cm^2$. The Bristle-spGal4-1 line is available from the Bloomington Stock Center (RRID: BDSC_71032).

---

[1] M. Kabra, A. Robie, M. Rivera-Alba, S. Branson, and K. Branson, "Jaaba: interactive machine learning for automatic annotation of animal behavior," *Nature Methods*, vol. 10, pp. 64–67, 2013.

[2] L. van der Maaten and G. Hinton, "Visualizing data using t-sne," *J. of Machine Learning Res.*, vol. 9, pp. 2579–2608, 2008.

### SUPPLEMENTARY FIGURES

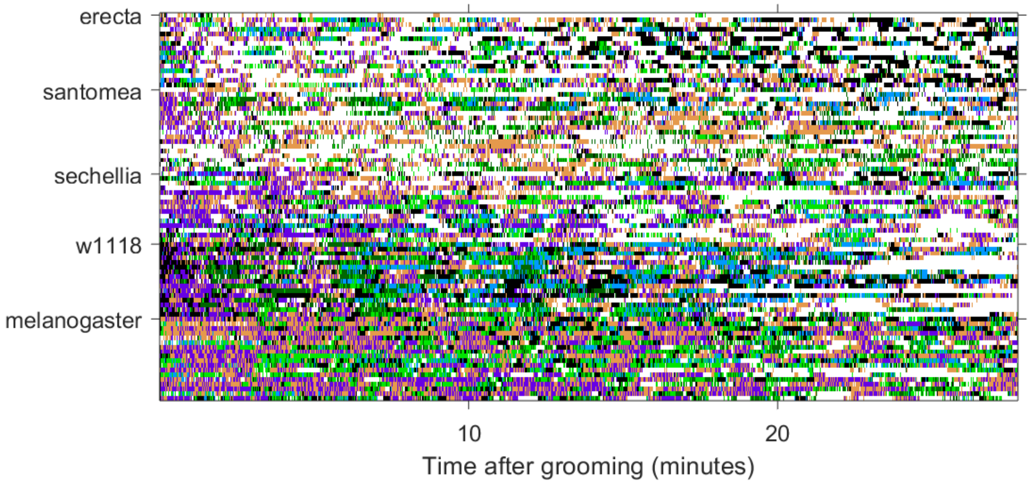


FIG. S1. **Ethogram for non-*melanogaster* drosophilids.** Shown are the ethograms for *erecta*, *santomea*, *sechellia*, and *simulans* flies. Rows are individual ethograms, columns indicate frame. Behaviors are coded by color. This is the full dataset associated with Figure 2.

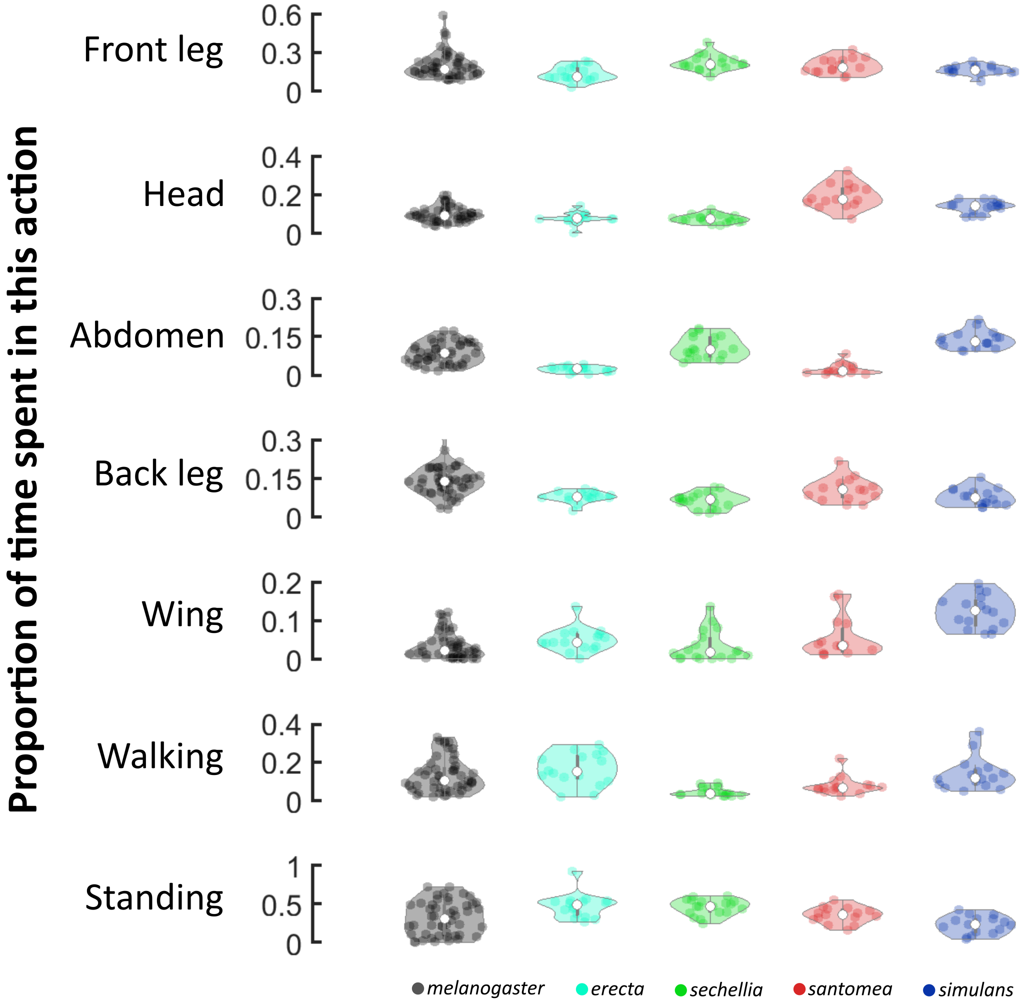


FIG. S2. **Behavioral proportions differ between species.** Shown are violin plots of behavioral proportions for five drosophilid species. Points indicate individual flies and color indicates species. Of the 70 pairwise comparisons between distributions, 25 (35.7%) possess different mean values (Kruskal-Wallis test, p<.05). 10 of the 25 differences are between *melanogaster* and non-*melanogaster* species, indicating that these species exhibit some differences in bulk behavior.

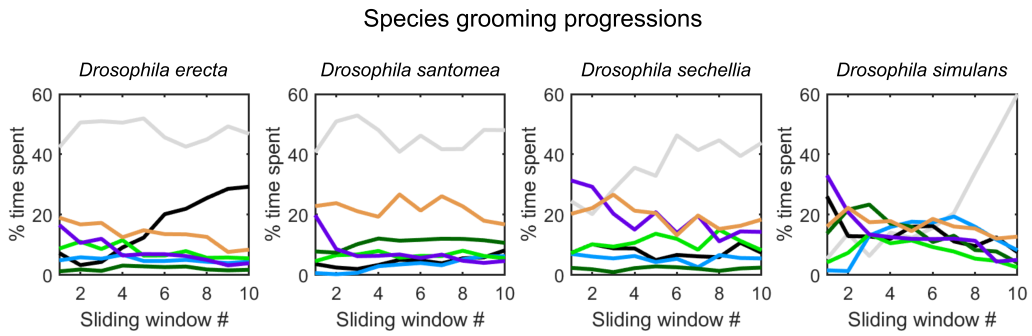


FIG. S3. **Grooming progressions are qualitatively similar across drosophilid species, with some noticeable differences.** Shown are the group mean grooming progressions for non-*melanogaster* populations. Grooming proportions were calculated across 10 non-overlapping windows (approximately 3 min. each). Y-axis indicates the proportion of time spent in an action. X-axis indicates the window. Note that all populations exhibit many similar trends, including an initial relatively high proportion of head grooming and front leg rubbing (purple and orange) followed by an increase in other grooming actions. Many populations differ significantly in their propensity to walk (black) or stand (gray) rather than groom. Though group means are shown here, individual vectors identical in format were used for classification analysis and visualization. These are the progressions associated with data in Figure 2.

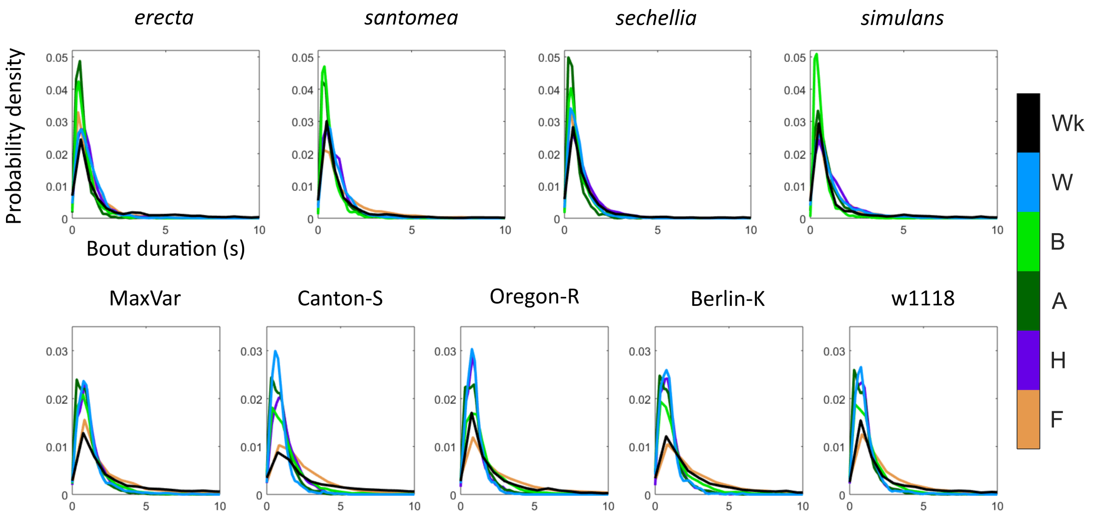


FIG. S4. **Bout duration distribution vectors of drosophilid species and *melanogaster* stocks** Shown here are the group mean bout duration distribution vectors for several populations. Y-axis indicates density. X-axis indicates bin. 20 bins of equal width were used, with the same bin width for all grooming actions. Non-grooming actions had larger bin widths due to longer average action durations for these actions. Note that non-*melanogaster* species perform a higher proportion of short actions, while *melanogaster* populations possess similar distributions to one another.

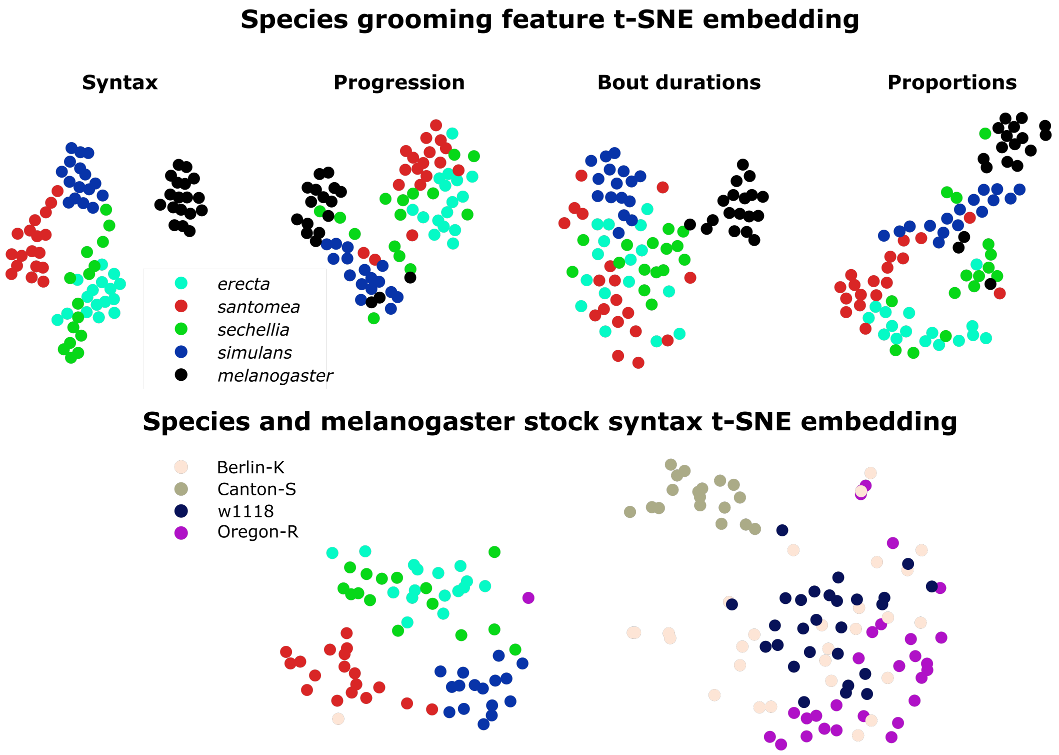


FIG. S5. **Grooming syntax separates species most strongly.** When comparing species, non-syntactic features of grooming do not exhibit the same degree of separability as grooming syntax. Shown here on top are t-SNE plots of each grooming figure displayed in Figure 1 of the main text, where points represent the grooming feature of an individual fly. Color indicates species. Qualitatively, a high degree of overlap between points indicates that grooming features are similar. Notably, grooming syntax provides the most distinct separation between species. However, other features also separate *melanogaster* from non-*melanogaster* species, suggesting that "bulk" or "average" grooming behavior is sufficient to distinguish between species. On the bottom is the t-SNE plot of syntax for non-*melanogaster* species and all *melanogaster* stock lines. Again, grooming syntax strongly separates *melanogaster* from non-*melanogaster* lines.

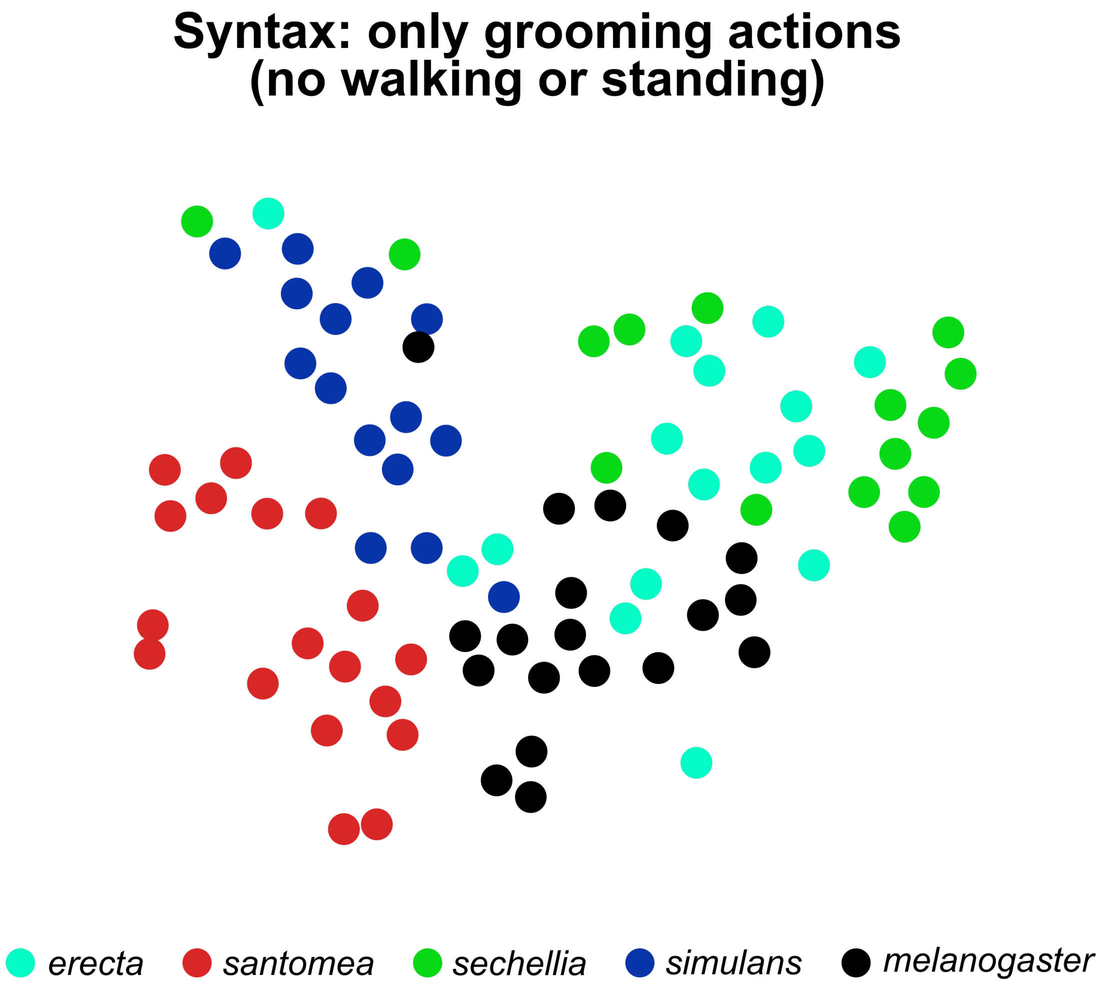


FIG. S6. **After removing walking and standing actions from syntax, species are still separable, but less so.** Shown here is a t-SNE embedding of species syntax with transitions to and from non-grooming actions (walking and standing) removed. When these non-grooming transitions are removed from syntax, individuals from different species resemble *melanogaster* (black) more closely (compare to Fig. S5 above for reference).

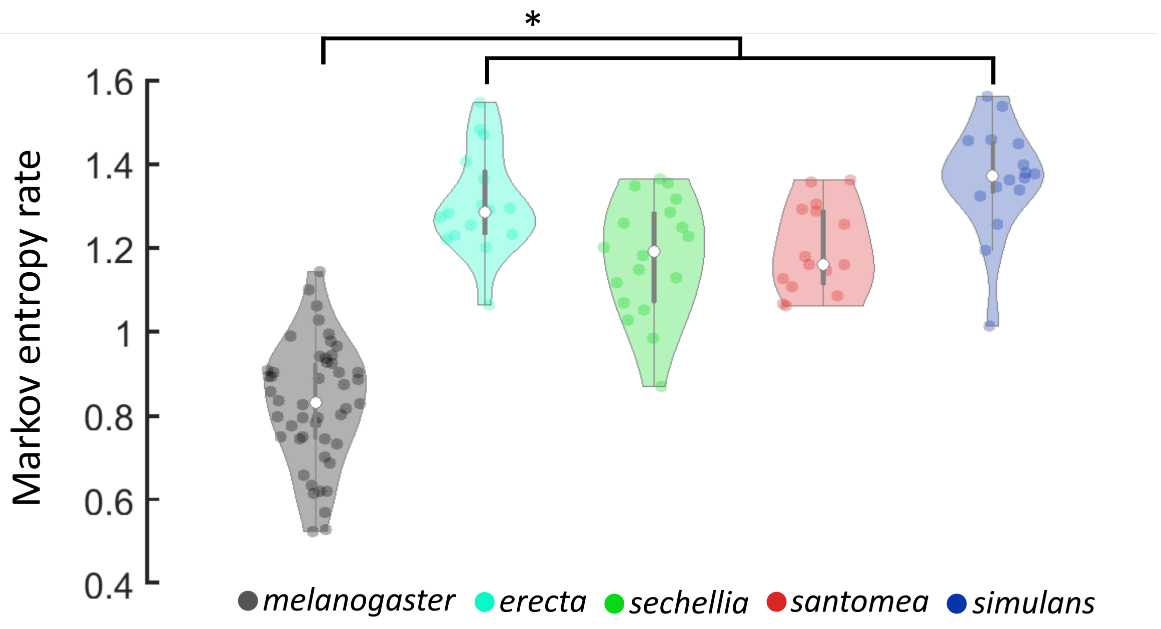


FIG. S7. ***melanogaster* grooming syntax is more stereotyped than non-*melanogaster* syntax.** Shown are violin plots of the Markov entropy rate of the grooming transition matrices calculated for each fly. Dots indicate individual flies and color indicates species. *melanogaster* flies exhibit slightly more stereotyped grooming transitions than non-*melaogaster* species, as indicated by the lower median entropy rate for *melanogaster* flies (Kruskal-Wallis test, p<.05). This increased stereotypy is driven by high anterior motif transition probabilities, which are illustrated in Figure 2B of the main text.

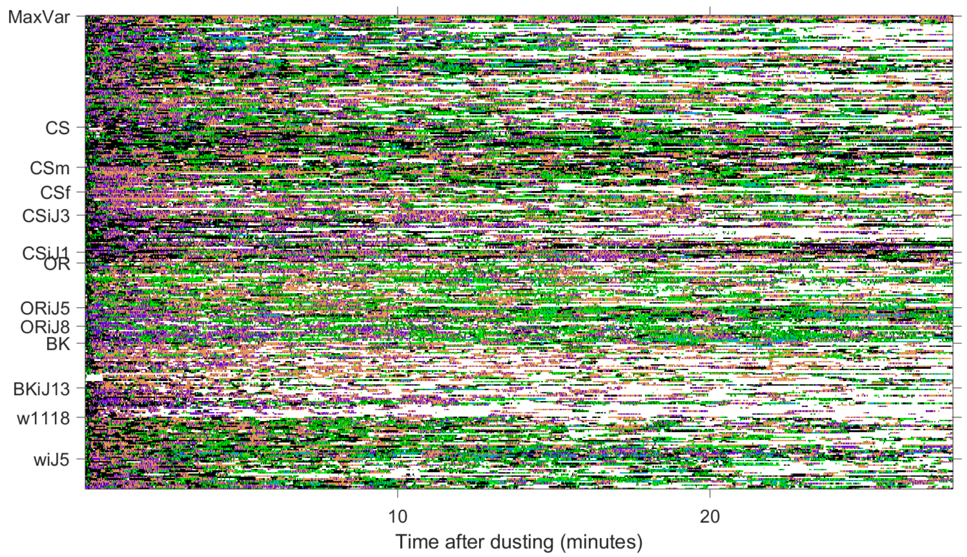


FIG. S8. **Ethogram for *melanogaster* flies.** Shown are the ethograms for each group of *melanogaster* flies analyzed here. Rows are individual ethograms, columns indicate frame. Behaviors are coded by color. This is the full dataset associated with Figure 3.

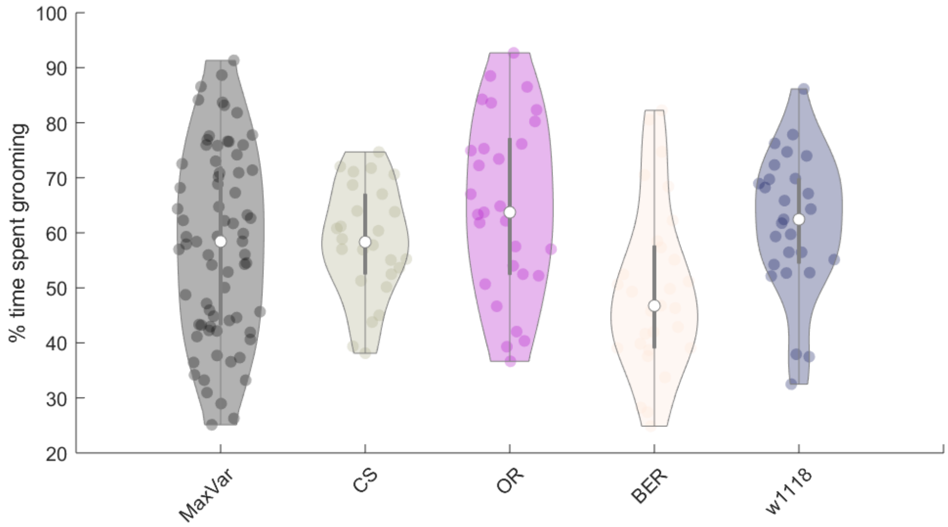


FIG. S9. **Different *melanogaster* populations exhibit robust grooming responses in this assay.** The amount of time spent grooming varies within species. However, regardless of genetic background, flies spend several minutes performing grooming actions after exposure to a dust stimulus. Shown are violin plots of grooming proportion distributions for five different *melanogaster* groups (right). White dots indicate median values. For all groups except Berlin-K, the median proportion of time spent grooming in a 27.8 minute recording after dusting is above 50% (13.9 minutes), indicating that nearly all flies exhibit a robust grooming response.

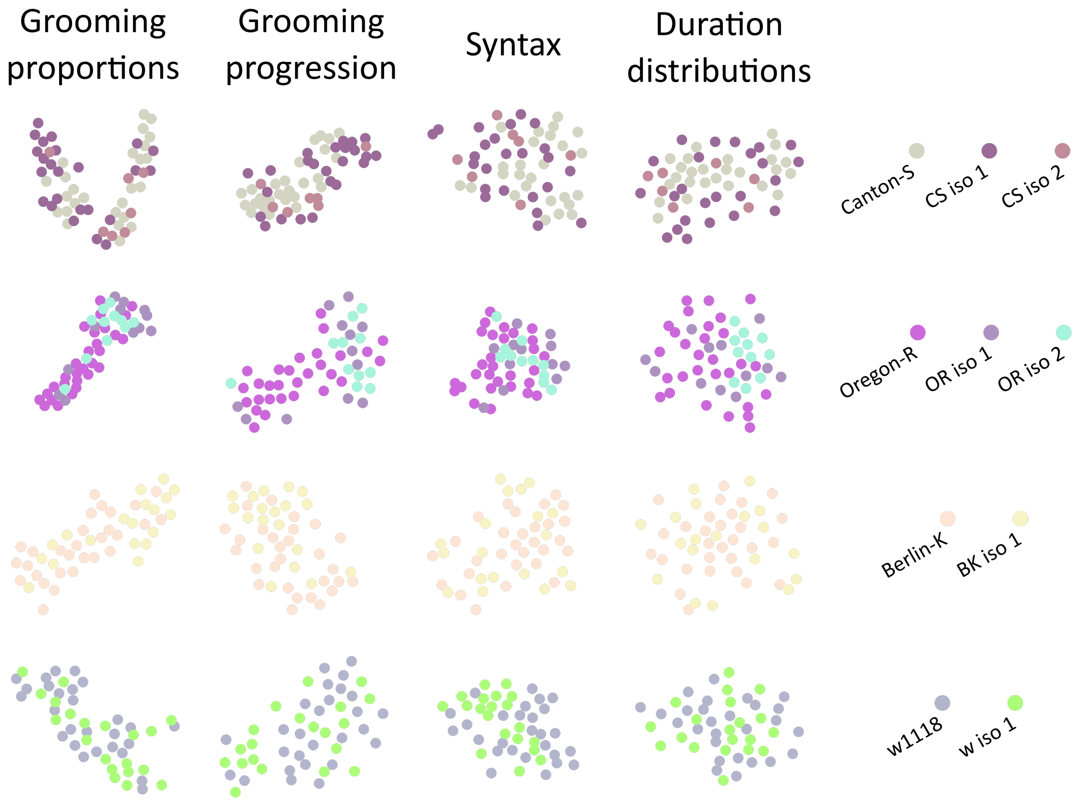


FIG. S10. **t-SNE of grooming features of *melanogaster* flies.** Shown are t-SNE embeddings of behavioral proportions, grooming progressions, grooming syntax, and bout duration distributions for all *melanogaster* stock lines and their derived isogenic strains. Each point indicates a feature for a single fly, and color (right) indicates group membership. The high degree of overlap by color indicates that isogenic strains do not differ markedly from their parent lines. However, the degree of "spread" or variability may be different between stock and isogenic lines, so variability in grooming syntax was explored in more detail in Figure 3 of the main text.

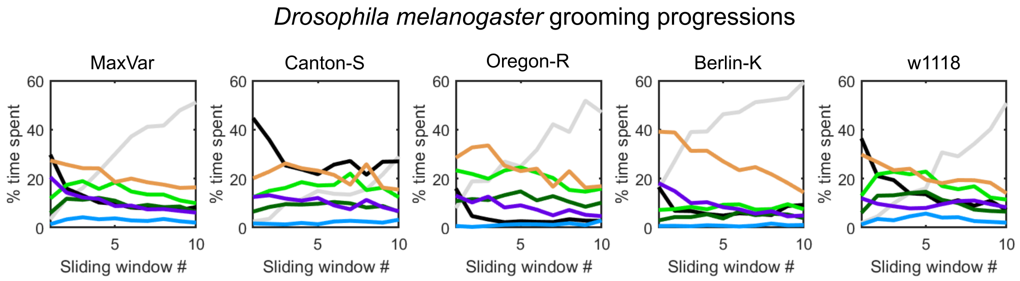


FIG. S11. ***Melanogaster* progressions differ in the amount of grooming and non-grooming actions.** Shown are the group mean grooming progressions for *melanogaster* populations. Grooming proportions were calculated across 10 non-overlapping windows (approximately 3 min. each). Y-axis indicates the proportion of time spent in an action. X-axis indicates the window. As with non-*melanogaster* species, all populations exhibited a general anterior to posterior grooming progression, with increased amounts of standing later in the progression. Canton-S flies tended to walk more than other stock lines, allowing for accurate classification when using grooming progressions as a feature. These are the progressions associated with data in Figure 3.

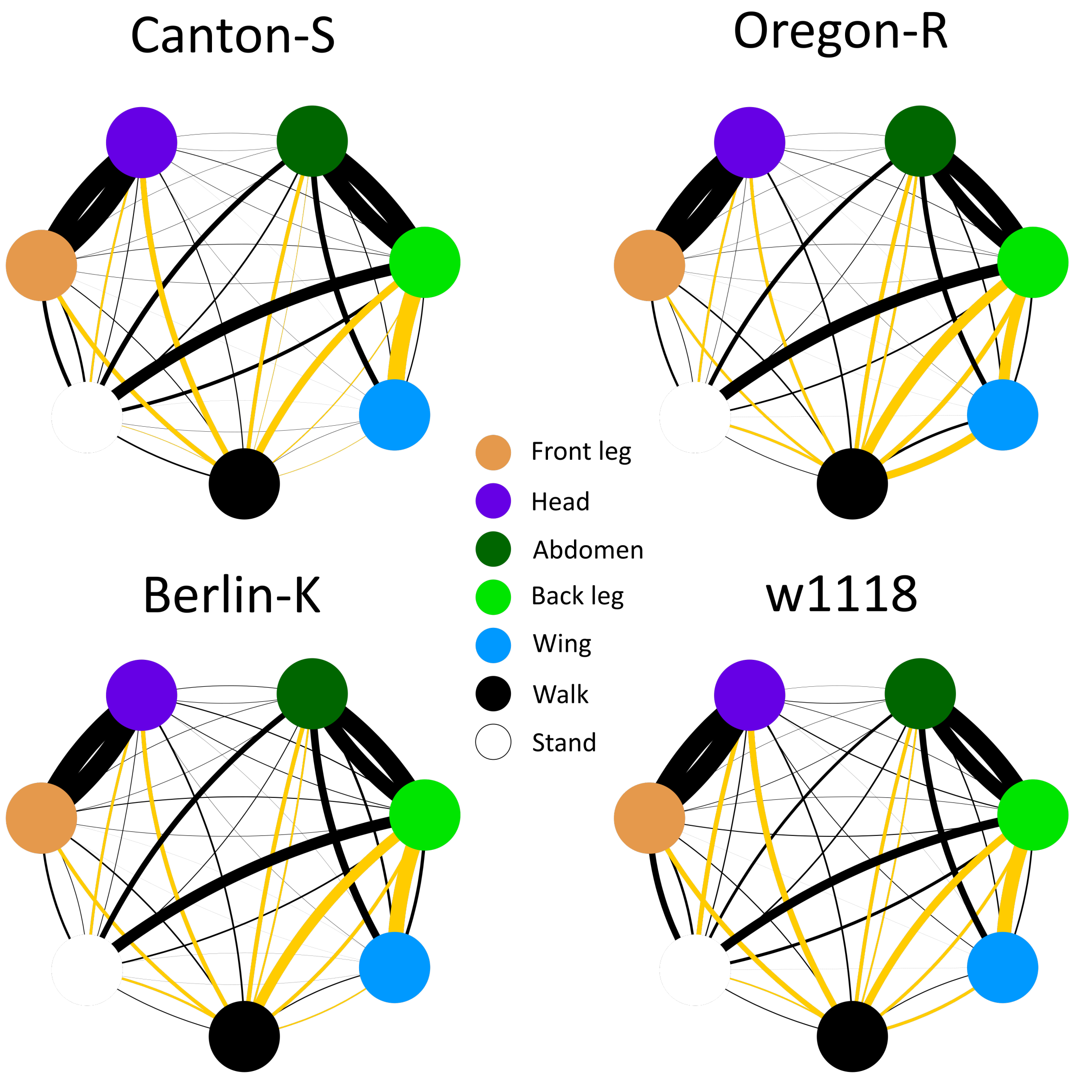


FIG. S12. **Grooming syntax of *melanogaster* stocks is highly conserved, but differences exist in transitions to and from non-grooming actions.** Shown are graphs of the group mean grooming syntax for each *melanogaster* stock. Nodes denote actions and edge thicknesses indicate transition probabilities between actions. Highlighted in gold are the top 10 transitions that differ most significantly between at least two stocks (Wilcoxon rank-sum test, $p < .05$ after Bonferroni correction). Only one of these transitions is within-motif (wing to back leg), indicating that grooming syntax is highly conserved within *melanogaster* stock lines. The other significantly different transitions involve walking or standing, indicating that stocks differ in overall activity levels.

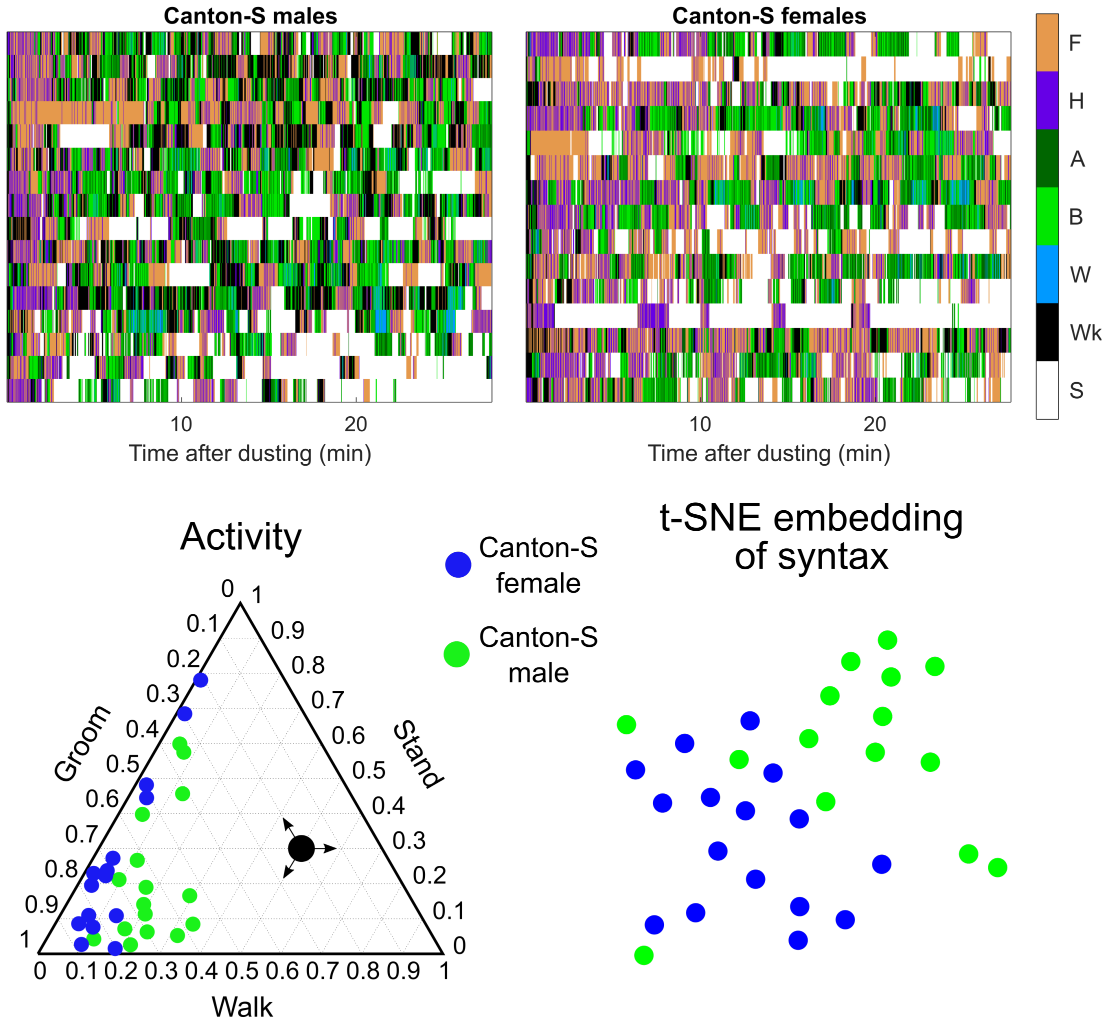


FIG. S13. **Male and female Canton-S flies possess distinct grooming features.** Shown on top are the ethograms for male and female Canton-S flies. Below is a ternary plot of the proportion of time spent grooming, walking, and standing for male and female flies. Female flies groom more than male flies, while male flies walk more than female flies. On the bottom right is a t-SNE embedding of male and female grooming syntax, with color indicating sex. The separation between sexes indicates that male and female flies possess distinct syntactic phenotypes. Logistic regression classification on grooming syntax confirms this, as it distinguishes between male and female flies with good accuracy (71%, chance levels would be 50%). Similar accuracy rates are achieved for progressions and grooming proportions, indicating that male and female flies possess identifiable differences in grooming behavior, largely driven by differences in locomotor behavior.

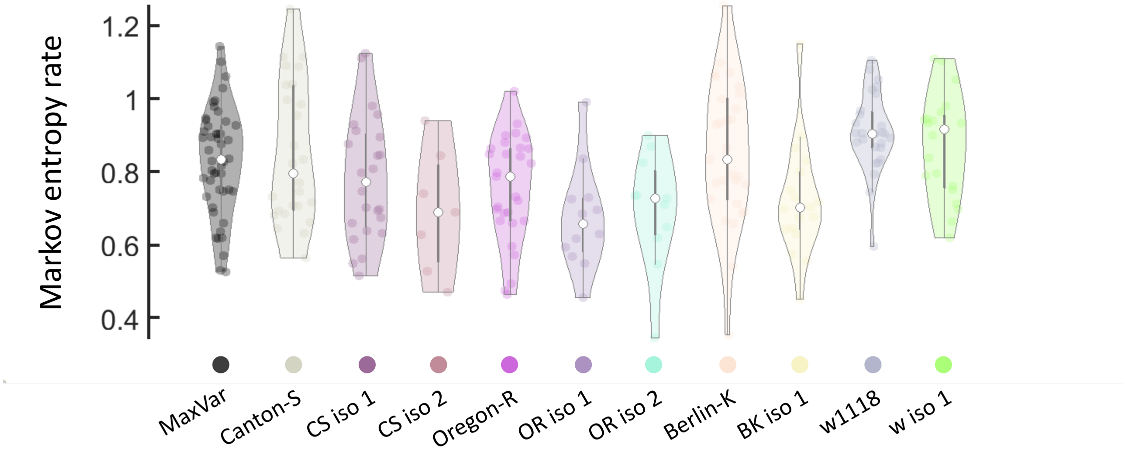


FIG. S14. ***melanogaster* lines exhibit similar degrees of grooming stereotypy.** Shown are violin plots of Markov entropy rates for *melanogaster* stock lines and isogenic strains. Dots indicate individual flies and color indicates group. None of the distributions shown here possess statistically significant medians (Kruskal-Wallis test, $p>.05$ for all pairwise comparisons), so all *melanogaster* strains perform grooming transitions that are similarly stereotyped.

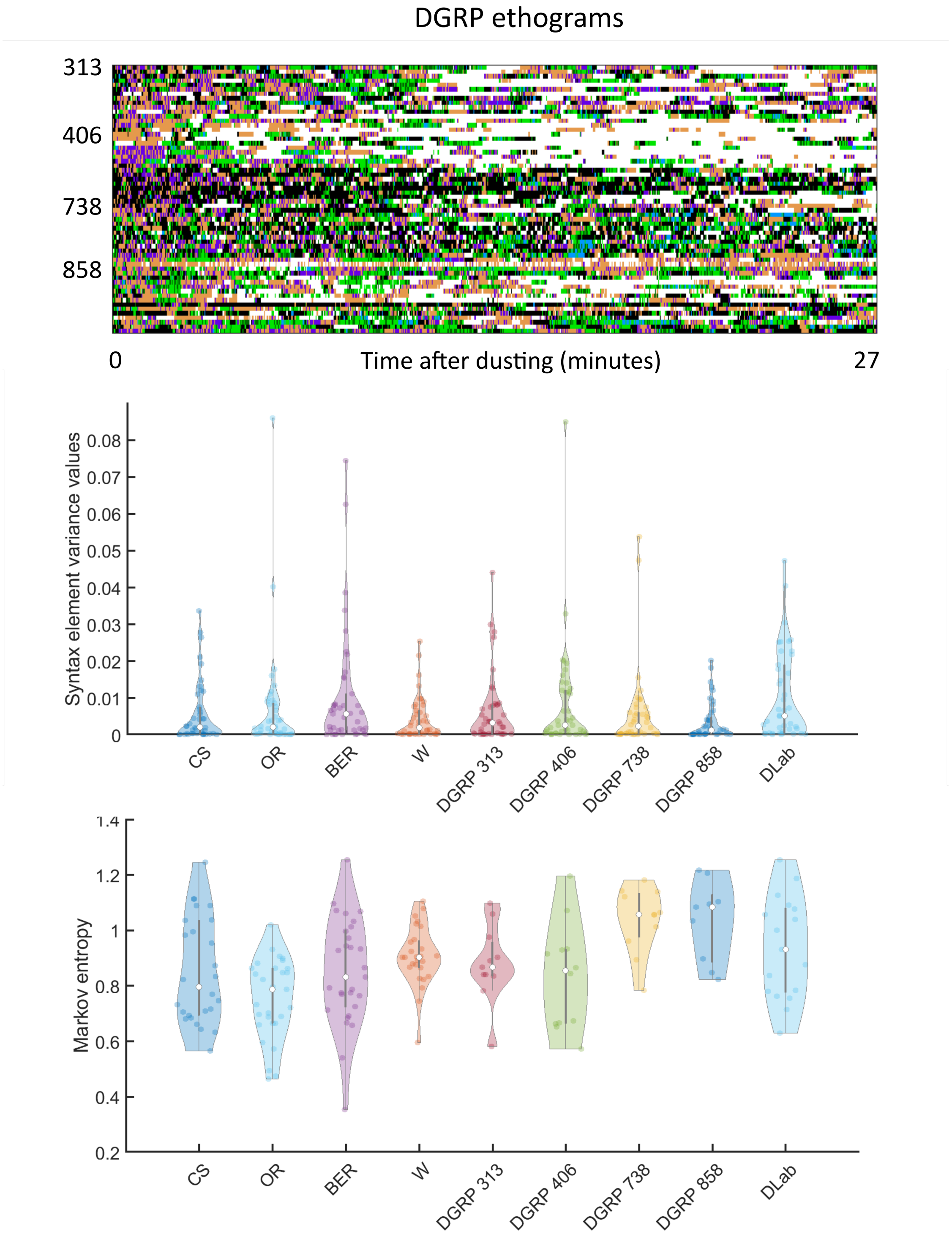


FIG. S15. **DGRP lines do not exhibit heightened grooming variability.** The syntax of four wild isolate DGRP lines and one line provided by the lab of Michael Dickinson (DLab) was compared to *melanogaster* stock lines. The top panel plots the variance values for each entry of the syntax matrix for each line. Wild-caught and DLab lines did not have significantly higher syntax element variance values than the stock lines analyzed in the main text. The bottom panel shows Markov entropy rates for each line. Again, DGRP and DLab lines were not less stereotyped (i.e., did not possess higher entropy rates) than stock lines.

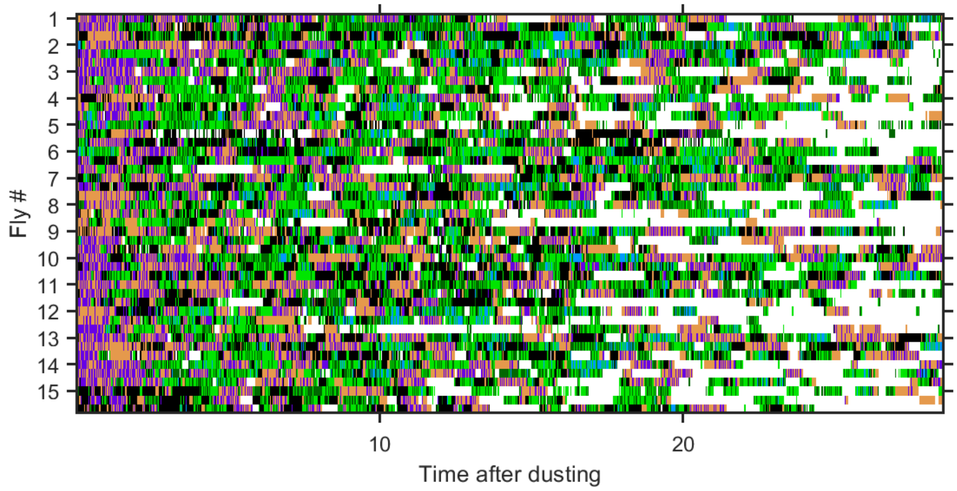


FIG. S16. **Ethogram for Canton-S individuals recorded across multiple sessions.** Shown are the ethograms for 15 Canton-S flies recorded for 3 sessions each over 3 consecutive days. Row labels indicate the individual fly, with rows after each label providing ethograms for sequential days (i.e. session 1, session 2, session 3). Columns indicate frame. Behaviors are coded by color. This is the full dataset associated with Figure 4.

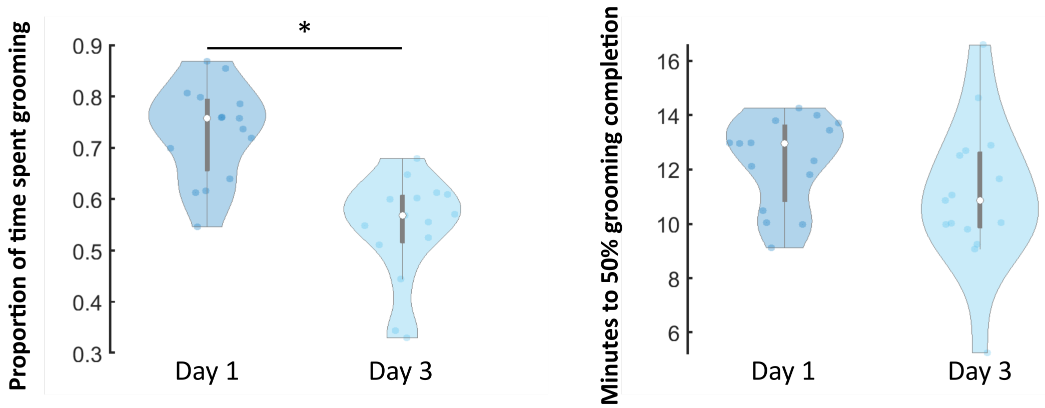


FIG. S17. **Individual flies groom less after several days of trials.** Shown on the left are violin plots of the proportion of grooming time (in a 30 minute session) of individual flies on day 1 compared to day 3 of experiments. Flies groom significantly less on day 3 of the experiment (paired t-test, p<.05). On the right are violin plots of the time a fly takes to complete half of their total amount of grooming for a session. These distributions do not differ, indicating that flies are not simply finishing their grooming sooner, but rather grooming in a more sporadic manner (i.e., walking and standing more between grooming bouts).

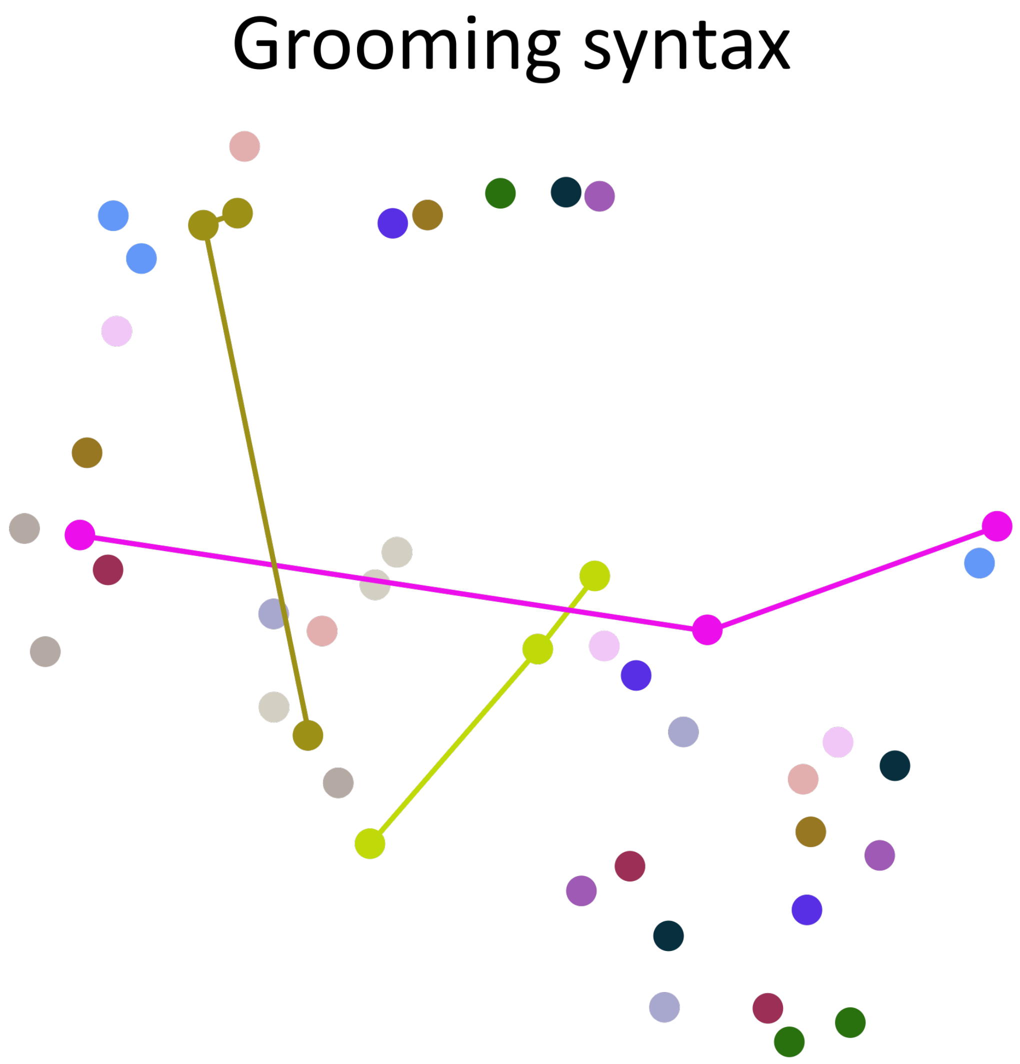


FIG. S18. **t-SNE of Canton-S individuals' grooming syntax recorded across multiple sessions.** Shown is the t-SNE embedding of the grooming syntax for 15 Canton-S flies recorded over 3 consecutive days. Dot color indicates fly identity. The absence of tight clusters of similarly colored dots indicates that flies do not resemble themselves more strongly over time than they do other flies. For illustration, three flies' syntax embeddings have been connected with colored lines; these three flies do not possess highly similar grooming syntax across 3 days, as indicated by the large distances between their syntax embeddings.

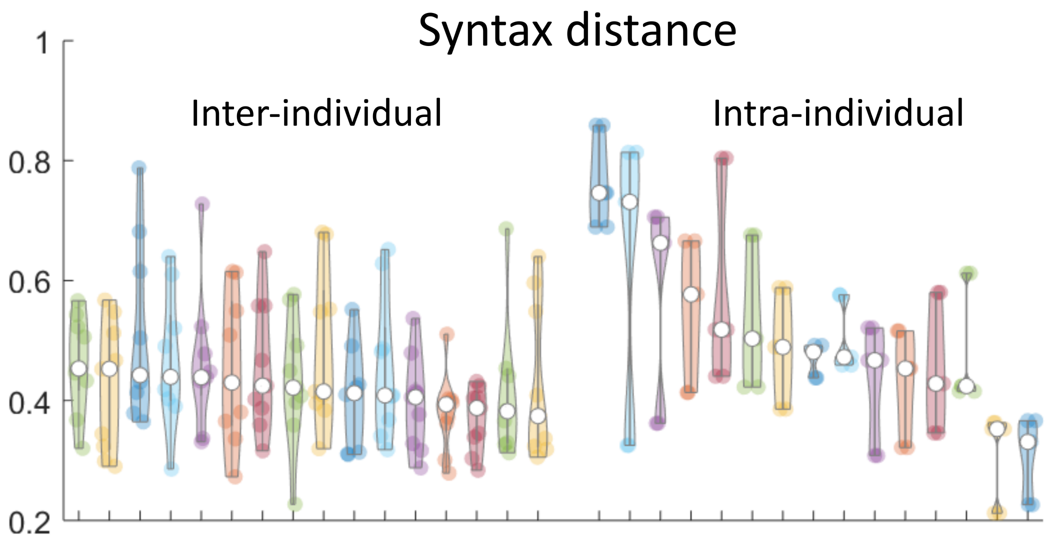


FIG. S19. **Canton-S individuals do not possess distinct grooming syntax.** 15 Canton-S flies were observed longitudinally three consecutive days (three recordings). Syntax for each fly was calculated for each session and pairwise Euclidean distances were calculated between syntax vectors. Shown here are violin plots of the distances between syntax vectors; points represent the distance between any two syntax vectors and hollow dots indicate the median. On the left are inter-individual comparisons, and on the right are intra-individual comparisons (i.e., comparing syntax vectors from the same individual on different days). The 15 most similar inter-individual comparisons are more similar than all of the intra-individual comparisons, indicating that flies do not possess strongly idiosyncratic grooming syntax over time.

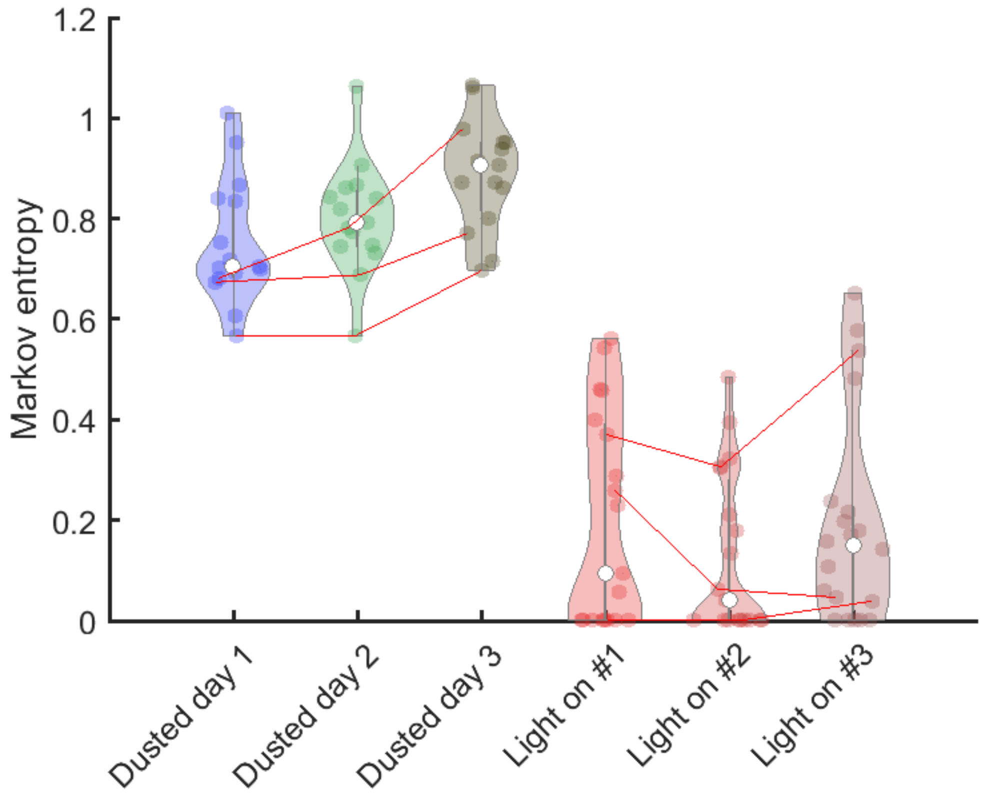


FIG. S20. **Sensory stimulus standardization reduces but does not abolish grooming variability.** Shown here are violin plots of the Markov entropy rates for dusted flies over three days and flies stimulated optogenetically for three sessions. Dots indicate individual flies and color indicates session. For comparison, see Figure 4C in the main text. For illustrative purposes, several examples of individual flies over several sessions have been connected using red lines (i.e., the lines connect the same individual over different sessions). Individual dusted flies exhibit grooming variability on a day-to-day basis, as illustrated by the changing Markov entropy rates. If individual syntax (or the degree of stereotypy) did not vary between days, these lines would be horizontal. Optogenetically-stimulated flies exhibit lower variability (as indicated by the lower entropy values), but can still exhibit session-to-session variability that is similar to dusted flies (indicated by the magnitude of changes in entropy between sessions). If flies exhibited perfect stereotypy, the entropy rate would be zero.

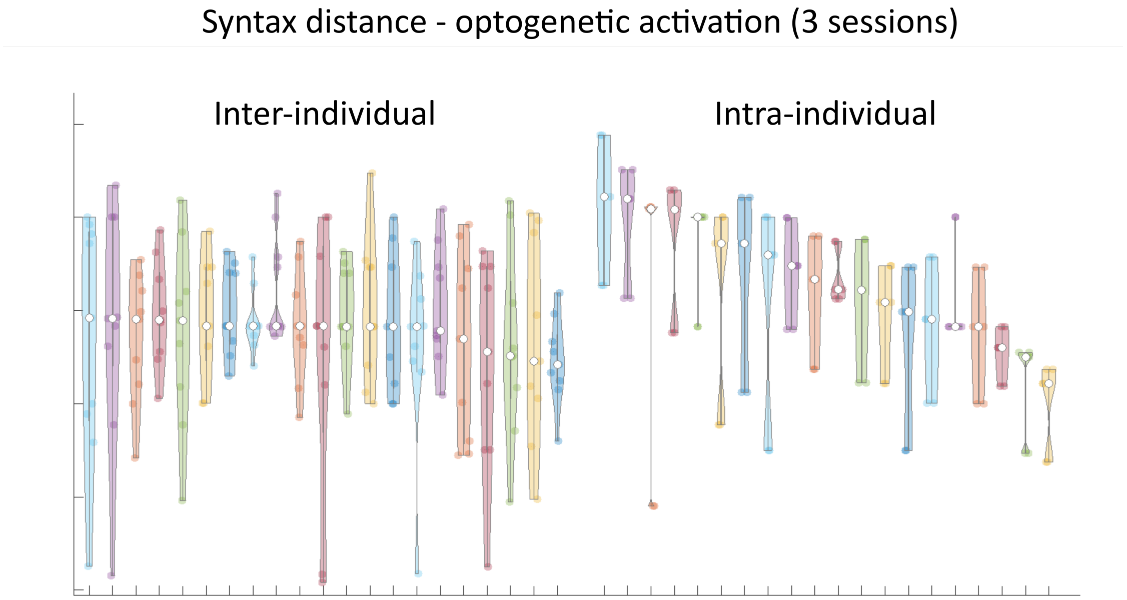


FIG. S21. **Optogenetic stimulation does not stereotype grooming syntax within individuals.** 20 flies were optogenetically stimulated to groom for three minutes over three sessions (see Figure 4 of the main text for experimental overview, Methods for further details). Syntax for each fly was calculated for each session and pairwise Euclidean distances were calculated between syntax vectors. Shown here are violin plots of the distances between syntax vectors; points represent the distance between any two syntax vectors and hollow dots indicate the median. Shown on the left are inter-individual comparisons, and on the right are intra-individual comparisons (i.e., syntax from the same fly compared over different optogenetic stimulation sessions). The 20 most similar inter-individual comparisons are as or more similar than all of the intra-individual comparisons, indicating that flies do not possess strongly idiosyncratic grooming syntax even when sensory input is standardized.
